# Vision-Based Genomic Model for Copy Number Variant Pathogenicity Prediction

**DOI:** 10.64898/2026.05.21.726953

**Authors:** Ilia Buralkin, Jorge Botas, Kai-Li Chang, Yueqian Deng, Athanasios Papastathopoulos-Katsaros, Zhandong Liu, Junseok Park

**Affiliations:** Graduate School of Biomedical Sciences, Baylor College of Medicine, Houston, TX, USA; Jan and Dan Duncan Neurological Research Institute, Texas Children’s Hospital, Houston, TX, USA; Department of Pediatrics-Neurology, Baylor College of Medicine, Houston, TX, USA

## Abstract

Copy number variants (CNVs) are a major class of structural genomic alterations underlying rare disease, including neurodevelopmental delay and intellectual disability, yet predicting their pathogenicity remains challenging. Existing methods reduce CNVs to region-level numerical features, discarding the positional structure and cross-track patterns that expert clinical reviewers use to interpret genomic evidence. To address this, we introduce Tesseract for CNV, a track-based spatial representation for CNV pathogenicity prediction, which represents each variant as a base-pair–resolution multi-track image and models spatial genomic patterns across annotation tracks while preserving positional structure and cross-track dependencies. Trained on a chromosome-level hold-out split of the ClinVar dataset, Tesseract outperforms prior methods on held-out and curated noncoding benchmarks, improving AUROC by up to 0.10 over the state-of-the-art baseline. On the independent DECIPHER cohort, the model demonstrates generalizability by maintaining the highest AUROC and the highest F1 score across baselines. Furthermore, our model localizes pathogenic signals to clinically meaningful genomic subregions, providing track-annotated evidence that supports practical clinical interpretation.

## 1 Introduction

Copy number variants (CNVs) are a class of structural variants involving deletions or duplications of genomic segments ≥ 50 bp, often spanning kilobases to megabases in length [1]. Unlike singlenucleotide variants (SNVs), CNVs can alter gene dosage and regulatory architecture across broad intervals, producing pathogenic effects through direct disruption of coding genes as well as indirect, position-dependent mechanisms such as enhancer–promoter disruption, topologically associating domain (TAD) boundary displacement, and cis-regulatory rewiring [2–4]. This ability to perturb both gene dosage and regulatory organization makes CNVs major contributors to rare genetic disease, including developmental delay, intellectual disability, neurodevelopmental disorders, and congenital anomalies [5–7]. However, because a single event can affect multiple genes and regulatory elements, many clinically relevant CNVs cannot be mapped to a single driver gene [8]. In clinical settings, pathogenicity is assessed under ACMG/ClinGen technical standards [9, 10], with expert review routinely relying on visual inspection of genomic and clinical annotation tracks in a shared spatial context (Figure 1A). This evidence-rich interpretation process already creates a scaling challenge for clinical screening, especially as genome-wide CNV pathogenicity interpretation continues to generate large numbers of candidate events requiring manual review. These pressures motivate automated tools that can classify CNV pathogenicity while preserving the multi-track spatial evidence structure used by expert reviewers.

**Figure 1:**
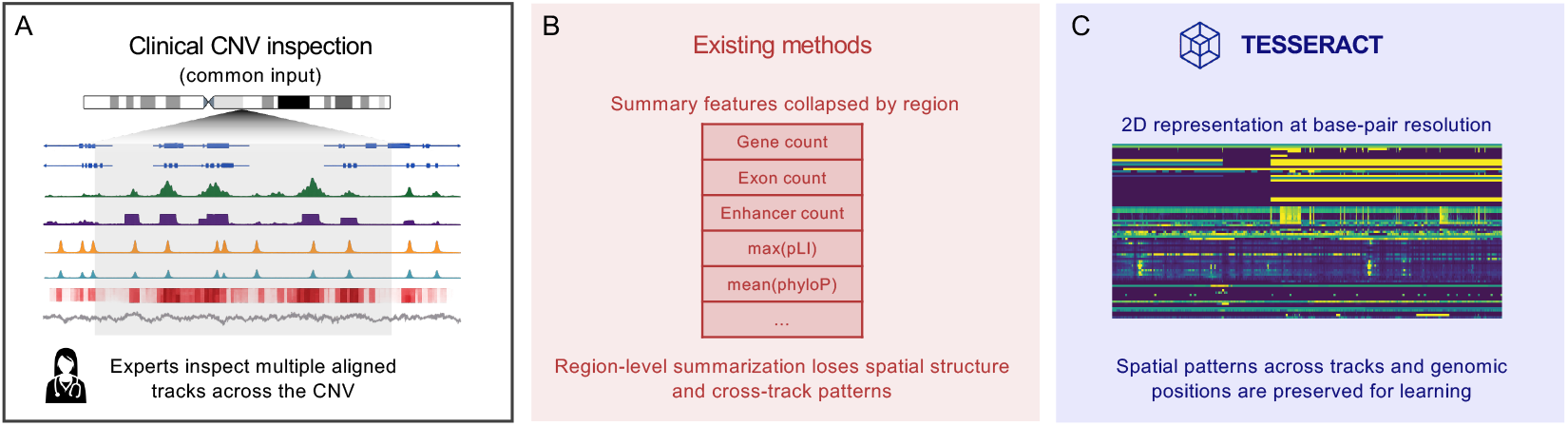
Tesseract preserves the multi-track spatial representation that clinicians use to interpret CNVs. **(A)** Clinical review of a CNV combines many coordinate-aligned annotation tracks (gene structure, conservation, regulatory marks, dosage labels) inspected jointly across the variant interval in a genome browser. **(B)** Prior CNV-pathogenicity tools collapse each track into a handful of scalar summary statistics over the variant interval, discarding both within-track positional structure and between-track co-occurrence before learning. **(C)** Tesseract instead represents each variant as a *T × W* multi-track image at base-pair resolution, with *T* = 78 annotation tracks along the height axis and *W* genomic positions along the width axis, and learns directly on this representation.

A range of computational tools have been developed to automate CNV pathogenicity prediction. Rule-based methods such as AnnotSV [11], SvAnna [12], ClassifyCNV [13], and AutoCNV [14] formalize ACMG/ClinGen-style criteria by assigning deterministic scores from overlapping annotations. Moving beyond rule-based methods, machine-learning methods including TADA [15], CNVoyant [16], and StrVCTVRE [17] train tree-based classifiers on engineered CNV-level summaries, such as gene counts, constraint scores, dosage annotations, and overlap statistics. More recent deep-learning approaches, such as PhenoSV [18], further increase modeling capacity by learning interactions among richer coding and noncoding interval-level features. Nevertheless, across existing methods, the input representation remains fundamentally region centric: each CNV is compressed into variant, gene, or interval summary features before learning (Figure 1B), rather than modeled directly from the underlying annotation landscape at base pair resolution. This compression discards the shared genomic coordinate system that clinical geneticists inspect visually across annotation tracks.

To address this representational gap, we introduce Tesseract, which interprets each CNV as a multi-track image at base-pair resolution, with genomic annotation tracks along one axis and genomic position along the other. A length-agnostic learning framework encodes local windows of this image and aggregates them across the CNV to produce a variant-level pathogenicity score for variants ranging from 50 bp to 1 Mbp. This representation preserves the coordinate-aligned, multi-track evidence that clinical geneticists use to visualize and interpret genomic evidence, rather than compressing it into region-level summary statistics before learning (Figure 1C). Across heldout ClinVar coding and curated noncoding benchmarks and the independent DECIPHER cohort, Tesseract achieves the strongest overall performance among evaluated tools, with the best AUROC and highest *F*_1_ score for pathogenicity classification. Its base-pair resolution representation further enables localization of pathogenic signal to specific CNV subregions, supporting clinical interpretation alongside variant-level prediction.

## 2 Related work

A major class of CNV pathogenicity prediction methods represents each variant using engineered CNV-level summary features and applies conventional tabular machine-learning models to these aggregated representations [17, 15, 16]. StrVCTVRE employs a random forest classifier trained on exonic structural variants, encoding each CNV using 17 handcrafted features that capture gene importance, conservation, coding-sequence structure, and expression-related properties of the affected region [17]. TADA adopts a similar feature-engineering framework, training separate deletion and duplication classifiers from fixed-length feature vectors derived from dosage-sensitive genes, known pathogenic intervals, and local regulatory or topologically associated domain (TAD) context [15].

CNVoyant further extends this paradigm to multi-class CNV classification using random-forest-based models trained on 17 curated CNV-level attributes, while additionally providing SHAP-based feature attribution explanations [16].

PhenoSV introduces a deep-learning-based formulation while preserving the same underlying representational assumption [18]. Each structural variant is decomposed into gene- or region-centric instances annotated with fixed-length feature vectors comprising variant impact scores (e.g. CADD), constraint metrics, conservation signals, and phenotype-aware statistics. These embeddings are processed using a transformer-based multiple-instance learning framework with masked multi-head attention to model both direct and indirect gene effects, followed by max pooling to produce variant-level and gene-level pathogenicity predictions [18]. Despite architectural differences, these methods all operate on pre-aggregated CNV- or region-level summaries rather than directly modeling basepair-resolution, coordinate-aligned multi-track genomic signals.

## 3 Methods

### 3.1 Data representation

#### From annotation tracks to a multi-track image

We represent each copy-number variant (CNV), specified by genomic coordinates (chrom, start, end), as a multi-track image **X** ∈ ℝ^*T* ×*W*^ at basepair resolution, with *T* = 78 genomic annotation tracks stacked along the height axis and *W* = (end − start) + 1000 positions along the width axis, where the extra 1000 bp provides 500 bp of flanking context on each side. This preserves the stacked, coordinate-aligned view of genomic annotations used in genome browsers such as Integrative Genomics Viewer(IGV) [19] and the UCSC Genome Browser [20], which support clinical CNV review (Figure 1A,C).

#### Input tracks

The 78 tracks span diverse classes of genomic annotations—gene structure and constraint, dosage sensitivity, evolutionary conservation, integrated SNV pathogenicity scores, regulatory and chromatin context, and ClinVar pathogenic SNVs and insertion or deletion (indel) positions— organized into 14 biological families, together with two variant-identity tracks (variant mask and log-normalized length) derived directly from the variant record. A detailed breakdown of the track groups, the rationale for excluding gnomAD structural-variant allele frequency [21], and per-track specifications are provided in Appendix A.1.1 and Appendix Table 1.

**Table 1:** The 78 input channels of Tesseract, organized into 14 biological families. Full per-track provenance, including source URLs and preprocessing notes, is provided as supplementary metadata.

| Group | # | Channels |
| --- | --- | --- |
| Variant identity | 2 | variant_mask, length_channel |
| Gene structure | 6 | GENCODE v48 (exon, CDS), lncRNA, 5'/3', UTRs, OMIM genes |
| Gene constraint | 7 | pHaplo, pTriplo, pLI, LOEUF, mutConstraint, CCRs, HI_index |
| Dosage sensitivity | 13 | clinGenGeneDisease + 12 HI/TS evidence-tier tracks |
| Conservation | 9 | phyloP (100/20/30-way), phastCons (100/20/30-way), GERP_RS, ConsHMM (50/100) |
| Integrated scores | 6 | LINSIGHT, CADD, CDTs, JARVIS, gwRVIS, GenoCanyon |
| Histone marks (ESC) | 9 | H2AFZ, H3K4me1/2, H3K9me3/ac, H3K27me3/ac, H3K36me3, H3K79me2 |
| Regulatory marks (ESC) | 6 | ATAC, DNase, CTCF, RAD21, POLR2A, SMC3 |
| Blood epigenome | 5 | ATAC (imputed), H3K27ac/H3K4me3 (observed), RNA ± strand |
| Regulatory elements | 9 | ChromHMM 15-state, super-enhancers, cCRE (5 classes), FIRE, ReMap CRM |
| Chromatin structure | 2 | TAD boundaries, 40 kb boundaries |
| Sequence context | 1 | GC content (5 bp) |
| Known pathogenic | 2 | ClinVar pathogenic SNVs, ClinVar pathogenic indels |
| Population frequency | 1 | gnomAD SNV allele frequency |
| **Total** | **78** |  |

#### Track preprocessing

Tracks are rendered at base-pair resolution under one of three regimes, selected per track. *Binary interval* sets **X**_*t,w*_ = 1 at every position covered by an annotation interval of track *t*, 0 elsewhere, with max-reduction on overlaps (e.g. GENCODE exons, ClinGen HI/TS evidence tiers). *Categorical with graded strength* renders ordered evidence tiers from four annotation sources under a fixed ordinal map to [0, 1] (e.g. the ChromHMM 15-state chromatin model). *Continuous normalization* applies a per-track transform *f*, clips to a per-track range 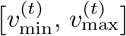 and linearly rescales to [0, 1],

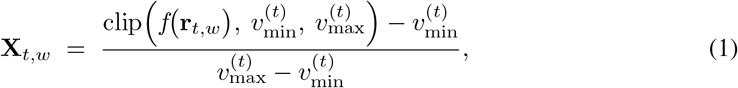

with *f* the identity or log(1 + *x*) depending on the track’s dynamic range (e.g. log(1 + *x*) for heavy-tailed blood histone-coverage tracks). gnomAD SNV allele frequency uses a dedicated sparse encoding that rescales −log_10_(AF) to [0, 1]. Full per-track regime assignments, transforms, and scaling parameters are listed in Appendix A.1.3.

### 3.2 Model architecture

Tesseract maps the variant image **X** ∈ ℝ^*T* ×*W*^ (Section 3.1) to a pathogenicity probability in four stages: a window encoder that produces per-window embeddings from non-overlapping 500 bp slices of **X**; an optional sequence model (Transformer[22] or Mamba-2[23]) that refines the resulting variable-length embedding sequence; a variant aggregator that collapses the sequence into a single representation **z**; and a classification head that predicts variant pathogenicity. Figure 2A shows the end-to-end pipeline (TESSERACT-E2E) where the window encoder is trained jointly with the downstream components on labeled CNVs. Figure 2B shows a self-supervised alternative where we pretrain the window encoder as a variational autoencoder on the unlabeled windows and freeze it before training the downstream components (TESSERACT-VAE).

**Figure 2:**
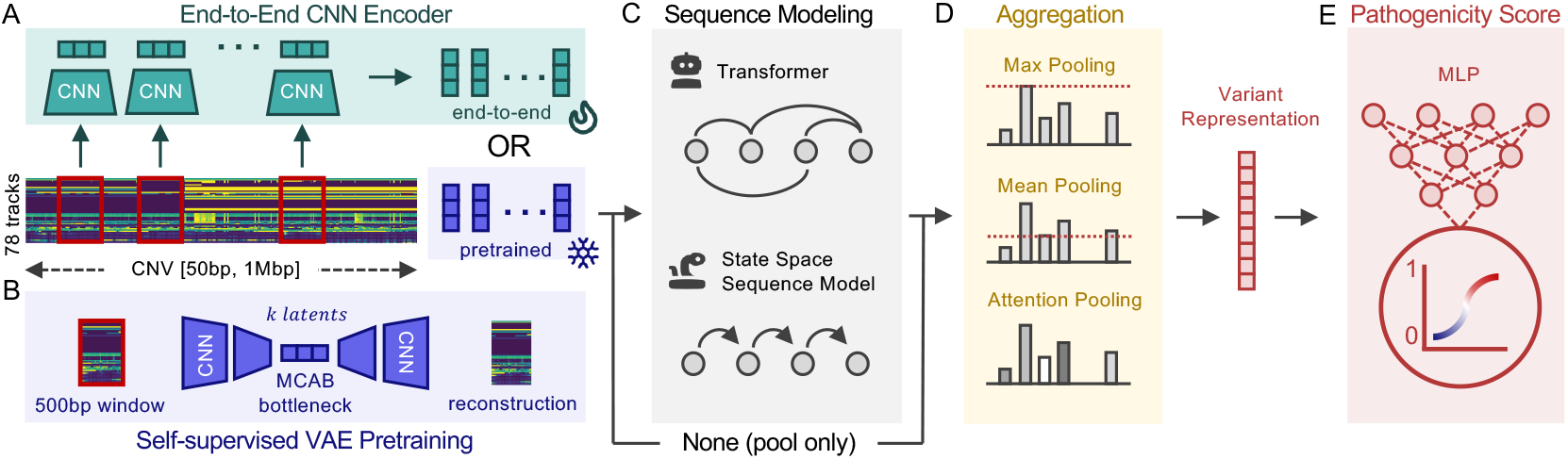
Tesseract architecture. The model maps a multi-track variant tensor to a pathogenicity score through four stages: a window encoder (**A** or **B**), an optional sequence model (**C**), a variant aggregator (**D**), and a classification head (**E**). **(A)** Tesseract-E2E uses an end-to-end 2D CNN encoder trained jointly with the downstream components on labeled CNVs. **(B)** Tesseract-VAE replaces the encoder with a frozen VAE pretrained on unlabeled 500 bp windows under a masked-reconstruction objective; downstream stages are identical between the two variants. **(C)** The resulting per-window embeddings are optionally refined by a Transformer or state-space (Mamba-2) sequence model, or passed through unchanged. **(D)** Embeddings are aggregated into a single variant-level representation by max, mean, or attention-based (ABMIL) pooling. **(E)** An MLP head maps the variant representation to the predicted pathogenicity probability.

**Figure 3:**
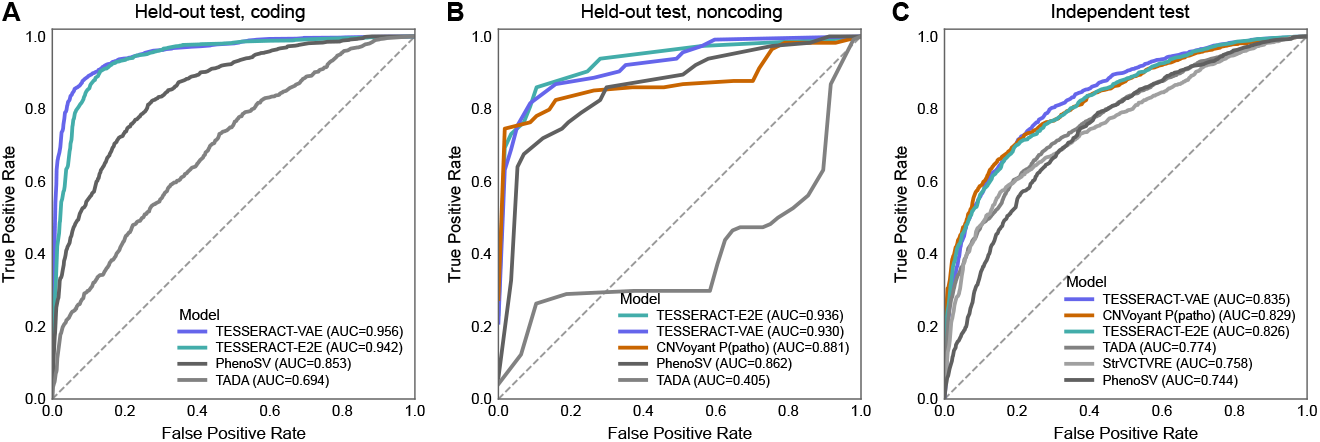
Main benchmark comparison on the ClinVar test split and the DECIPHER independent test cohort. Panels A–C compare Tesseract against published baselines on the in-distribution coding split, the matched in-distribution noncoding split, and the out-of-distribution DECIPHER cohort, respectively.

#### CNN window encoder

Each 500 bp window is encoded by a compact two-dimensional convolutional network whose kernels span both the track axis (height) and the genomic-position axis (width). The encoder follows the MobileNetV2 inverted-bottleneck design [24] with squeeze-and-excitation [25], a combination chosen for parameter efficiency at per-window encoding. Four stages of inverted-residual blocks produce a fixed-dimensional embedding **h**_*k*_ for a given 500bp window.

#### VAE window encoder

To leverage the substantially larger pool of unlabeled 500 bp windows relative to labeled CNVs, we pretrain a variational autoencoder [26, 27] under a masked-reconstruction objective [28], then freeze its encoder for downstream classification. The VAE reuses the same CNN tokenizer described above and compresses each window into 64 latent tokens using a cross-attention bottleneck [29]. For downstream classification, the latent tokens are mean-pooled into a single embedding **h**_*k*_ ∈ ℝ^128^. Full details are provided in Appendix A.2.1.

#### Sequence backbone

The window embeddings 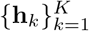 are optionally refined by either no sequence model, a 4-layer Transformer encoder [22], or a 4-layer Mamba-2 state-space model [30]. The identity setting ablates sequence modeling entirely, while the Transformer and Mamba-2 variants allow information exchange across genomic windows before aggregation. The three options define one axis of the ablation grid evaluated in Section 4.2.2.

#### Variant aggregator

Because CNVs range from 50 bp to the megabase scale, each variant is represented by a variable number of 500 bp windows. The aggregator is therefore designed to accept variable-length sequences and produce a fixed-dimensional variant-level representation. Given the sequence 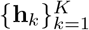, the aggregator collapses it to a single representation **z**. Standard aggregation stratigies include masked mean and max pooling over windows; we instead use attention-based multiple-instance learning (ABMIL) [31], which assigns each window *k* a gated attention weight

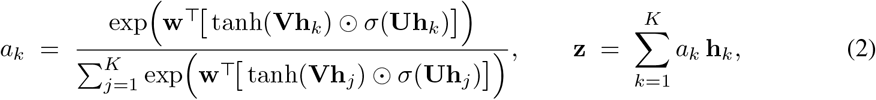

with learnable parameters **w, V, U** and *σ* the logistic sigmoid. ABMIL produces a data-dependent soft selection over windows; the per-window weights {*a*_*k*_} are retained alongside the variant-level score and used as an interpretability signal in Section 4.3.2. We compare ABMIL against mean and max pooling in the ablation study of Section 4.2.2. The pooled representation **z** is then passed through a two-layer MLP classification head to produce the variant-level pathogenicity probability.

#### Primary TESSERACT model configurations

Our full ablation grid spans two window-encoder regimes (end-to-end convolutional, frozen VAE), three sequence backbones (none, Transformer, Mamba-2), and three pooling operators (ABMIL, mean, max); the complete grid is presented in Figure 4 (Section 4.2.2). Two configurations serve as the primary models in the main-text comparison: Tesseract-E2E, an end-to-end convolutional architecture without an explicit sequence backbone and with ABMIL pooling; and Tesseract-VAE, which pairs a frozen pretrained VAE encoder with a 4-layer Transformer backbone and ABMIL pooling.

**Figure 4:**
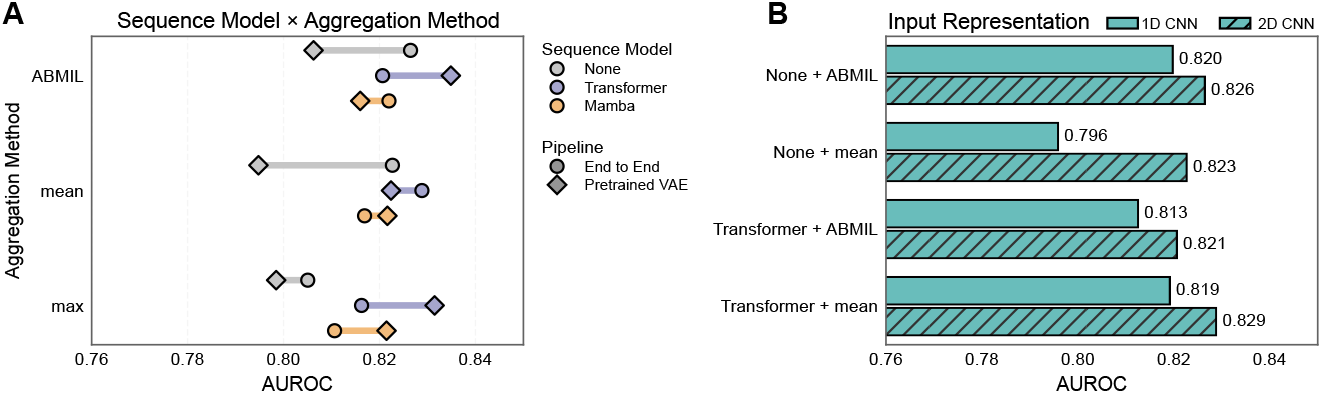
Architecture ablations for Tesseract. Panel A sweeps encoder regime (circle: end-to-end, Tesseract-E2E; diamond: pretrained VAE, Tesseract-VAE), sequence backbone (colour), and pooling operator (rows) on independent test at segmented inference with *S* = 120. Panel B compares the 2D channel-as-height representation against a 1D per-channel alternative for the strongest Tesseract-E2E configurations.

### 3.3 Datasets and splits

#### Training data

We train and internally validate Tesseract on the CNV dataset constructed by PhenoSV [18], using its original chromosome-level train/validation/test split. After preprocessing, the dataset contains 19,477 deletion and duplication CNVs (9,229 benign; 10,248 pathogenic), with coding CNVs drawn from ClinVar[32] and matched noncoding CNVs assembled from the PhenoSV[18] source datasets. This split evaluates held-out generalization to unseen chromosomes, reduces train/test leakage, and enables direct comparison to PhenoSV[18]. Source datasets, preprocessing details, coding/noncoding definitions, and per-split statistics are provided in Appendix A.3 and Table 2.

**Table 2:**
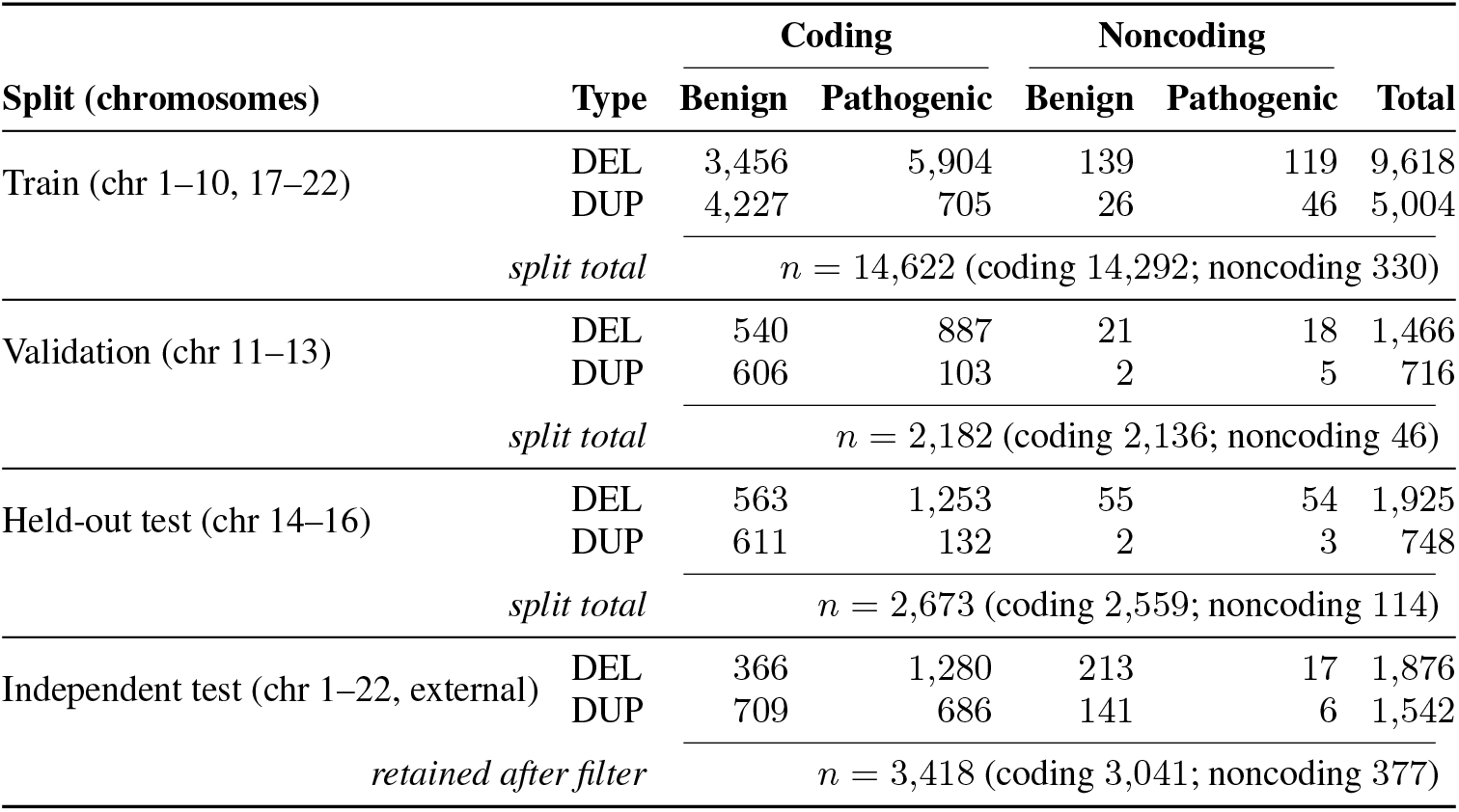
Per-split CNV counts by label, SV type, and coding/noncoding status. TESSERACT-CNV is trained on the chromosome-stratified corpus of PhenoSV [18], partitioned identically. Chromosomes appear in exactly one split each. The Independent test (DECIPHER-derived) is an external evaluation set (Section 3.3) scored without retraining; its row counts are reported after the reciprocaloverlap leakage filter and the restriction to the DEL/DUP classes scored by all baselines (rationale in Appendix A.3.2).

| Split (chromosomes) | Type | Coding |  | Noncoding |  | Total |
| --- | --- | --- | --- | --- | --- | --- |
|  |  | Benign | Pathogenic | Benign | Pathogenic |  |
| Train (chr 1–10, 17–22) | DEL | 3,456 | 5,904 | 139 | 119 | 9,618 |
|  | DUP | 4,227 | 705 | 26 | 46 | 5,004 |
| | *split total* | $n = 14,622$ (coding 14,292; noncoding 330) | | | | |
| Validation (chr 11–13) | DEL | 540 | 887 | 21 | 18 | 1,466 |
|  | DUP | 606 | 103 | 2 | 5 | 716 |
| | *split total* | $n = 2,182$ (coding 2,136; noncoding 46) | | | | |
| Held-out test (chr 14–16) | DEL | 563 | 1,253 | 55 | 54 | 1,925 |
|  | DUP | 611 | 132 | 2 | 3 | 748 |
| | *split total* | $n = 2,673$ (coding 2,559; noncoding 114) | | | | |
| Independent test (chr 1–22, external) | DEL | 366 | 1,280 | 213 | 17 | 1,876 |
|  | DUP | 709 | 686 | 141 | 6 | 1,542 |
| | *retained after filter* | $n = 3,418$ (coding 3,041; noncoding 377) | | | | |

**Table 3:** Independent test breakdown. Counts after the four-step preprocessing pipeline applied to the DECIPHER v11.15 release, restricted to coding variants (at least one base pair of overlap with a GENCODE v48 protein-coding exon). Class balance differs across SV types: deletions are predominantly pathogenic, whereas duplications are roughly balanced, reflecting the underlying distributions of clinically submitted DECIPHER records rather than a choice of the authors.

| SV type | Benign | Pathogenic | Total |
| --- | --- | --- | --- |
| DEL | 366 | 1,280 | 1,646 |
| DUP | 709 | 686 | 1,395 |
| Total | 1,075 | 1,966 | 3,041 |

#### Independent test

For external evaluation under distribution shift, we construct an independent test cohort from DECIPHER v11.15 [33], excluded from all training and validation procedures. Following the preprocessing protocol of PhenoSV[18], we additionally apply reciprocal-overlap filtering against the training corpus (90% for same-label overlaps, 85% for cross-label overlaps) to remove potential duplicate or label-conflicting events, and restrict evaluation to coding deletions and duplications. The final cohort contains 3,041 coding variants and is used consistently across all benchmark comparisons. Additional preprocessing details are reported in Appendix A.3.2.

### 3.4 Training and inference

#### Objective and optimizer

All configurations are trained to minimize binary cross-entropy on the variant-level logit using AdamW [34] with cosine learning-rate decay and early stopping on validation AUROC. The self-supervised VAE pretraining stage uses a separate optimization recipe; full hyperparameters for both supervised training and pretraining are provided in Appendix A.4.1.

#### Segment-based window sampling

CNV lengths vary substantially, resulting in a highly nonuniform number of 500 bp windows per variant. To construct fixed-size training inputs while maintaining coverage across the full event, we adopt the segment-based sparse sampling strategy of Temporal Segment Networks [35], partitioning each variant tensor into *K* = 120 equal-length segments and sampling one window uniformly from each segment at every training step. Resampling windows at every epoch additionally acts as a spatial augmentation. Unless otherwise noted, we use *K* = 120, matching the default inference segment size *S* = 120.

#### Annotation-group dropout

To prevent the model from collapsing onto a small subset of dominant tracks, we apply group-wise dropout over the 14 annotation families during training, independently zeroing each group with fixed probability. For the self-supervised configuration, we instead apply 0.1 dropout directly to the frozen VAE embeddings before sequence modeling.

#### Inference

Tesseract supports two inference modes for variable-length CNVs. In *full-length* inference, the complete window sequence is processed in a single forward pass. In *segmented* inference, the sequence is partitioned into fixed-size segments of length *S*, each segment is scored independently, and segment-level predictions are aggregated into a variant-level score. Unless otherwise noted, segmented inference uses `max` aggregation, with top-*k* mean (the mean of the *k* highest per-segment probabilities) evaluated as an alternative aggregation strategy (Section 4.3.1). Main-text results use segmented inference at *S* = 120 with `max`; full-length counterparts and segment-size sweeps are reported in the supplementary appendix and Figure 5, respectively.

**Figure 5:**
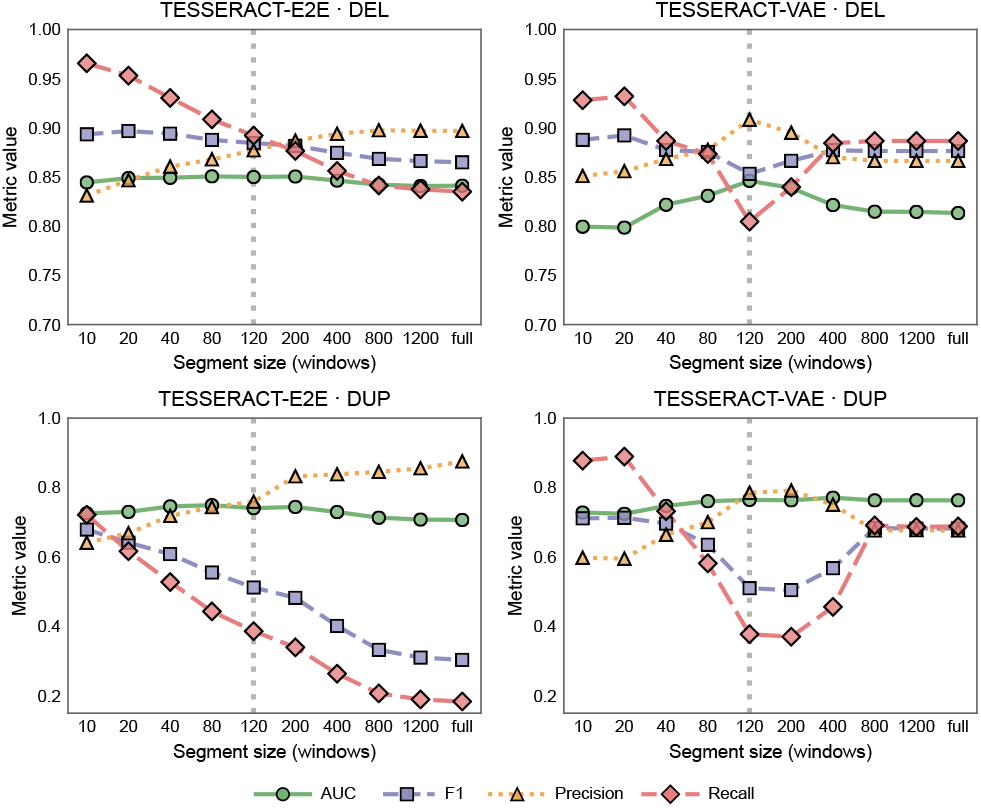
Segment-size sweep on independent test (*n* = 3,041). Rows = SV type (top: deletions; bottom: duplications), columns = model (left: Tesseract-E2E; right: Tesseract-VAE). Each panel plots AUROC, *F*_1_, precision, and recall at threshold 0.5 as a function of inference segment size *S* ∈ {10, 20, 40, 80, 120, 200, 400, 800, 1200} windows under `max` reduction over per-segment probabilities. Dashed vertical reference at the training-matched *S* = 120.

## 4 Experiments

### 4.1 Experimental setup

We evaluate Tesseract on two benchmarks that together probe in-distribution and out-of-distribution generalisation. The *held-out test* evaluates in-distribution performance on chromosomes excluded from training under the PhenoSV chromosome-level split protocol [18]. The noncoding subset uses the matched benchmark design from the same split. The *DECIPHER independent test set* [33] serves as the out-of-distribution benchmark. Unless otherwise noted, the main benchmarking results report the two primary configurations introduced in Section 3.2: Tesseract-E2E and Tesseract-VAE. The rationale for these selections, together with the full architecture sweep, is presented in the ablation study of Section 4.2.2.

We compare Tesseract against four published CNV-pathogenicity tools: CNVoyant [16], PhenoSV [18], TADA [15], and StrVCTVRE [17]. Each baseline is evaluated only on data outside its training set; split-specific reporting constraints and the rationale for including or excluding each baseline on a given benchmark are summarised in Appendix A.4.2. PhenoSV, which is trained on the same data as Tesseract, is reported on every evaluation split as an in-distribution reference point.

#### Evaluation metrics

We report three metric families on every benchmark: AUROC and average precision (AP) for ranking, Brier score on *P* (pathogenic) for calibration, and *F*_1_, precision, and recall at the operating point. Operating-point metrics use a fixed decision threshold of 0.5 for all probabilistic scorers and the native argmax over {benign, Variant of Uncertain Significance (VUS), pathogenic}for CNVoyant’s three-class softmax.

### 4.2 Main results

#### 4.2.1 Benchmarking against state-of-the-art

Tesseract is the strongest overall model family in the main benchmark comparison (Figure 3): Tesseract-VAE is the strongest ranker across both in-distribution splits and the external cohort, whereas Tesseract-E2E is often stronger at the default operating point, particularly on noncoding variants and under distribution shift.

##### In-distribution performance

On ClinVar held-out test coding (*n* = 2,559; Figure 3A), Tesseract-VAE leads on ranking (AUROC 0.956, AP 0.965), followed by Tesseract-E2E (0.942 / 0.947), both ahead of PhenoSV (0.853 / 0.869) and TADA (0.694 / 0.734); under fulllength inference both retain strong operating-point performance (Tesseract-E2E *F*_1_ = 0.900; Tesseract-VAE *F*_1_ = 0.866). The matched noncoding split (*n* = 114; Figure 3B) shows the same ranking—Tesseract-E2E at 0.936 / 0.950 and Tesseract-VAE at 0.930 / 0.943 ahead of CNVoyant (0.885 / 0.927) and PhenoSV (0.862 / 0.865). Although CNVoyant ranks noncoding variants well, its native three-class argmax assigns all scored variants to BEN, collapsing pathogenic-class F1 to zero (Table 6); Tesseract-E2E maintains the strongest thresholded performance, reaching F1 = 0.827 versus 0.659 for Tesseract-VAE. Full metrics are in Tables 4 and 5.

##### Out-of-distribution generalization

On independent test (*n* = 3,041; Figure 3C), Tesseract-VAE achieves the highest AUROC (0.835, AP 0.899), with CNVoyant (0.829 / 0.906) and Tesseract-E2E (0.826 / 0.901) close in ranking and StrVCTVRE and PhenoSV trailing. Ranking similarity does not translate to comparable operating-point behaviour: at threshold 0.5, Tesseract-E2E reaches *F*_1_ = 0.778 and Tesseract-VAE 0.752, whereas CNVoyant collapses to 0.468 despite its strong AUROC, indicating that its three-class softmax is poorly calibrated for binary pathogenicity decisions out-of-distribution. Under full-length inference, Tesseract-VAE reaches the best observed *F*_1_ on this split (0.809) at a modest AUROC cost (0.815; full metrics in Table 7). The next subsection examines the architectural trade-offs underlying the two configurations’ complementary strengths.

#### 4.2.2 Architecture ablations

Figure 4 isolates the design choices behind the two headline configurations. The two encoder regimes have opposite backbone requirements (Figure 4 A): end-to-end training does not benefit from sequence modeling, whereas the frozen VAE encoder becomes competitive only when paired with a Transformer. Within the Tesseract-E2E family, the 2D channel-as-height representation additionally outperforms the 1D alternative in every paired comparison (Figure 4 B).

On Tesseract-E2E, sequence modeling adds little: at *S* = 120 the nine configurations span a narrow AUROC range of 0.805–0.829, with Transformer+mean reaching the top AUROC (0.829), no-backbone+ABMIL reaching the top AP (0.901), and no-backbone+mean within 0.003 AUROC of the top cell, so the simpler no-backbone setting is the stronger default. The frozen VAE encoder shows the opposite pattern. Without a backbone, the best VAE configuration reaches AUROC 0.806; adding a 4-layer Transformer lifts the VAE models into the 0.822–0.835 range, with Tesseract-VAE (Transformer+ABMIL) as the top cell at 0.835. The same configuration also achieves the highest *F*_1_ on this split under full-length inference (0.809; §4.2.1), consistent with frozen reconstruction-trained latents needing downstream inter-window refinement that an end-to-end encoder learns implicitly. Both patterns persist under full-length inference; full numbers for all 18 grid cells are in Table 8.

The 2D channel-as-height layout outperforms the 1D per-channel alternative across all four paired Tesseract-E2E configurations tested. At *S* = 120, the AUROC gain ranges from +0.007 to +0.027, largest for no-backbone+mean, supporting the view that clinically relevant CNV evidence is carried by cross-track patterns rather than within-track signals alone (full paired results in Table 9). The training-time window count *K* is also a tuned choice: retraining Tesseract-VAE at *K* ∈ {80, 120, 200, 400, 1000} confirms *K* = 120 as the most robust setting, matching the best peak performance in the sweep with the smallest full-length degradation (−0.020 AUROC versus −0.026 to −0.030 for other values; Table 10).

### 4.3 Analysis

#### 4.3.1 Segmented vs full-length inference

Inference segment size *S* is a deployment-time tuning axis rather than a single optimum. Deletions are ranking-stable for both Tesseract configurations across the sweep (Figure 5), but duplications separate the two models: Tesseract-E2E drifts toward conservative calling, with recall falling from 0.72 at *S* = 10 to 0.19 at *S* = 1200 and *F*_1_ from 0.68 to 0.31, while rising precision keeps AUROC stable at 0.71–0.75. In contrast, Tesseract-VAE-DUP traces a U-shape, with recall bottoming at 0.38 at *S* = 120 and recovering to 0.69 at *S* = 800. This trade-off yields three useful Tesseract-VAE deployment regimes: **ranking** (*S* = 120/`max`, AUROC 0.835), **screening** (*S* = 20/`max`, *F*_1_ = 0.822, recall 0.917), and **precision** (*S* = 120/top-10 mean, precision 0.925). The aggregator is a second-order tuning axis at fixed *S*: `max` remains the *F*_1_-optimal default at every *S*, but at fine *S*, top-*k* mean increases precision at the cost of recall (e.g. at *S* = 20, *P* = 0.75 → 0.83, *R* = 0.92 → 0.72 for `max` vs top-20 mean; Table 12). Full numerics by model × SV × *S* are in Table 11.

Duplications are harder than deletions for every classifier on this cohort, but Tesseract-VAE is among the narrowest-gap tools on the split. At *S* = 120, Tesseract-E2E and Tesseract-VAE show DEL–DUP AUROC gaps of +0.110 and +0.082 respectively, while compared baselines span +0.099 (CNVoyant) to +0.136 (TADA) (Appendix 13). This asymmetry reflects the greater difficulty of duplication pathogenicity prediction, driven by context-dependent copy-gain effects and weaker representation in curated pathogenic CNV datasets.

#### 4.3.2 Pathogenicity signal localization

Beyond a variant-level score, Tesser-act produces two spatially aligned localisation maps that together identify the genomic subregion driving each call. At fine-*S* inference (*S* = 10–20; 2.5–5 kb per segment), where Tesseract-VAE achieves its strongest operating-point performance on independent test (§4.3.1), the per-segment probability vector is a localisation readout: high-probability segments mark the subregion the model deems pathogenic. Full-length ABMIL attention provides a second, spatially matched map of the windows the classifier up-weights during aggregation. Figure 6 illustrates these views on a pathogenic *TCF4* deletion (DECIPHER 259631, 69 kb). At segmented inference with *S* = 10, the predicted probability exceeds 0.5 over a contiguous block centered on the gene body. The strongest full-length ABMIL attention peaks fall on the same region alongside positive dosage-sensitive and disease-gene annotations, while benign-direction tracks remain inactive.

**Figure 6:**
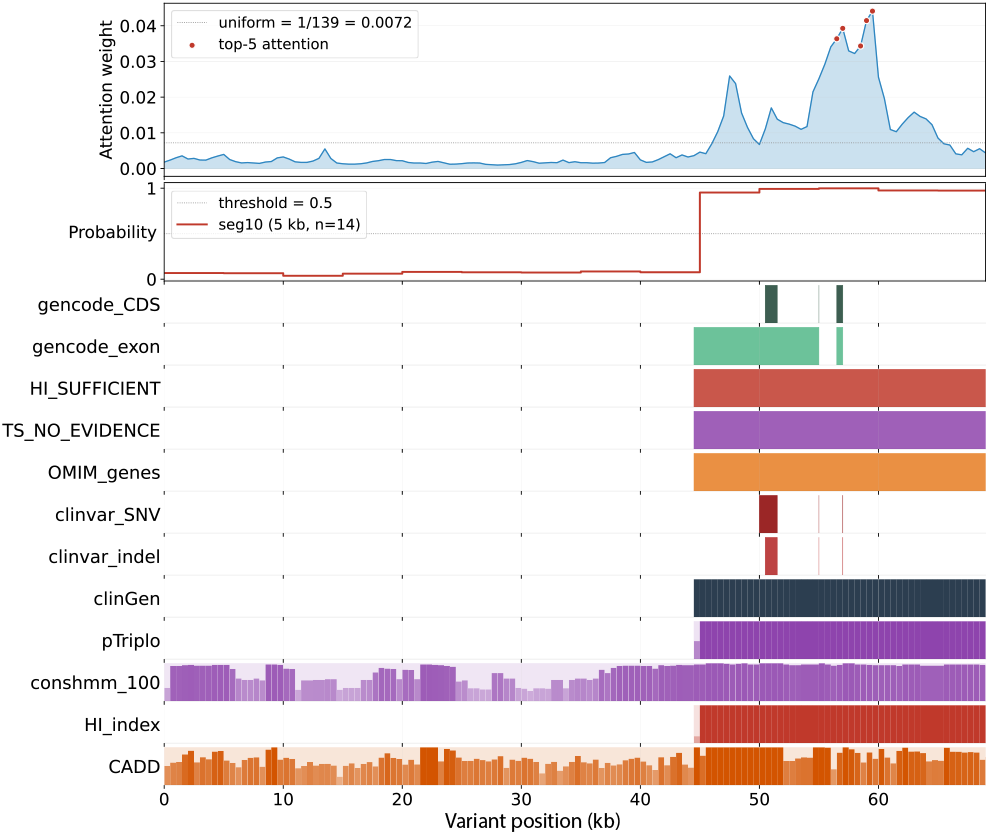
CNV signal localization on a pathogenic *TCF4* deletion (DECIPHER 259631, chr18:55,177,922–55,246,617, 69 kb, *n* = 139 windows). Top: full-length AB-MIL attention. Middle: per-segment probabilities at *S* = 10. Bottom: subset of aligned annotation tracks supporting the call (full per-channel statistics in Appendix 14).

The within-variant alignment seen on *TCF4* reproduces at cohort scale. Across CNVs in the independent test, curated dosage- and disease-gene annotations rank as the strongest correlates of per-segment probability, with clinGenGeneDisease the single strongest channel (*ρ* = +0.47), followed by conservation tracks and then weaker epigenetic marks (Appendix 14). Benign-direction channels (gwrvis, CDTS, HI_UNLIKELY) remain significantly negative, providing an internal null that the model shifts probability away from the no-evidence side of the dosage annotations.

Together, these readouts provide complementary localization views across genomic scales: full-length ABMIL attention highlights the windows most strongly weighted during aggregation, while segmented inference localizes pathogenic signal to progressively smaller subregions as S decreases. Across both the case study and cohort-level analyses, the strongest localization signals consistently align with dosage-sensitive, disease-gene, and conservation annotations. This pattern is concordant with the supplementary channel-perturbation analysis, where curated disease-gene, dosage-related, length, and known pathogenic SNV/INDEL tracks produced larger performance changes than most broad regulatory, epigenomic, or chromatin groups (§A.10). Because the VAE encoder is pretrained with masked reconstruction and the end-to-end model is trained with annotation-group dropout, these perturbations reflect post-training evidence reliance rather than a direct estimate of each track group’s intrinsic biological importance. Taken together, the localization and perturbation analyses suggest that Tesseract learns a clinically meaningful spatial representation of CNV evidence, while also motivating a more careful distinction between clinical evidence use and mechanistic biological modeling.

## 5 Conclusion and Discussion

TESSERACT improves CNV pathogenicity prediction by reframing clinical annotations as a spatial representation-learning problem rather than a region-level feature-engineering problem. Instead of summarizing each CNV into aggregate scores, TESSERACT preserves the coordinate-aligned structure of genes, constrained elements, dosage-sensitive regions, regulatory annotations, and prior pathogenic evidence across the affected interval. This design better matches clinical CNV review and provides the strongest overall utility among evaluated tools across both in-distribution and out-of-distribution benchmarks. The localization analyses further show that fine-segment inference identifies high-scoring subregions aligned with biologically plausible evidence, while full-length ABMIL attention provides a complementary view of the windows most influential for the variant-level prediction. Together, these readouts position TESSERACT not only as a pathogenicity classifier, but also as a spatial evidence-prioritization framework for CNV interpretation.

At the same time, the perturbation analyses clarify which evidence streams contribute most strongly to TESSERACT predictions. As expected for a model trained on limited clinical CNV labels, predictions are most sensitive to curated disease-gene, dosage-related, length, and known pathogenic SNV/indel tracks, while broader regulatory, epigenomic, and chromatin groups often have smaller individual effects. This reflects a broader limitation of supervised CNV pathogenicity modeling, where labels are shaped by the same clinical evidence used during expert interpretation. Future work could move beyond this annotation-dependent regime by combining TESSERACT’s spatial framework with long-context genomic foundation-model representations, such as AlphaGenome[36] or Evo2[37], to jointly model sequence context, gene regulatory network, evolutionary constraint, and dosage mechanisms not fully captured by available genomic tracks. Such hybrid models could preserve clinical interpretability while improving generalization to poorly characterized regions and supporting more objective pathogenicity assessments of de novo CNVs.

## A Appendix

This appendix collects implementation details that are useful for reproducibility but would interrupt the flow of the main paper. Subsections A.1–A.4 mirror the main-text methods section (§3); subsequent subsections back the figure panels reported in Section 4.

### A.1 Data representation

#### A.1.1 Input tracks: design rationale

##### Variant-identity tracks

Two tracks are derived directly from the variant record rather than from external annotation files. The *variant mask* takes value 0.5 inside the CNV interval for deletions and 1.0 for duplications, with 0 in the 500 bp flanking padding, jointly encoding variant type and the variant/flank boundary. The *length channel* holds a log-normalized CNV length (log svlen − log 50)/(log 10^6^ − log 50), clipped to [0, 1] and filled across the variant span (0 in padding); the effective length range is therefore 50 bp to 1 Mb.

##### Excluded input: gnomAD structural-variant allele frequency

We deliberately exclude gnomAD structural-variant allele frequency [21] from the input track set to prevent label leakage, since low allele frequency for structural variants is expected to inversely correlate with their functional significance [38]. We retain only gnomAD single-nucleotide-variant allele frequency, which is scored over a disjoint variant class and does not share this leakage risk.

#### A.1.2 Track groups

Full per-track provenance, including primary data source and download URL, is provided in the supplementary metadata file. The compact grouping in Table 1 is intended as a readable summary of the 78-channel input representation.

#### A.1.3 Track preprocessing (full specification)

##### Binary interval-fill

Tracks stored as genomic intervals (BED/BigBed format) without a continuous score column are binarized: we set **X**_*c,t*_ = 1 for every genomic position *t* that overlaps at least one annotation interval in channel *c*, and **X**_*c,t*_ = 0 otherwise, using max-reduction when intervals overlap. This regime applies to gene-structure masks from GENCODE v48, including exons, coding sequences (CDS), long non-coding RNAs (lncRNAs), and 5^′^/3^′^ untranslated regions (UTRs); OMIM disease-gene annotations; ClinVar pathogenic single-nucleotide variant (SNV) and insertion/deletion (indel) markers; ENCODE candidate cis-regulatory element (cCRE) classes, including promoter-like signatures (PLS), proximal and distal enhancer-like signatures (pELS and dELS), CTCF-only elements, and DNase-H3K4me3 elements; topologically associating domain (TAD) boundaries; and ClinGen haploinsufficiency/triplosensitivity (HI/TS) tracks.

##### Continuous normalization

BigWig continuous signal tracks—phyloP (100/20/30-way) [39], phastCons (100/20/30-way) [40], GERP_RS [41], CADD [42], LINSIGHT [43], GenoCanyon, JARVIS, gwRVIS, CDTS, `HI_index`, CCRs, FIRE, ReMap CRM, GC content, histone-mark coverage in ESC (H2AFZ, H3K4me1/2, H3K9ac/me3, H3K27ac/me3, H3K36me3, H3K79me2) and in Blood (imputed ATAC), transcription-factor / cofactor ChIP (CTCF, RAD21, POLR2A, SMC3), chromatin-accessibility assays (ATAC-seq, DNase-seq), and the ENCODE cCRE aggregate score— are read at base-pair resolution, transformed by a per-track function *f*, clipped to [*v*_min_, *v*_max_], and rescaled linearly to [0, 1] under Equation 1 in main text. Across the 39 BigWig-backed continuous tracks, 34 use *f* = identity (raw percentile-clipped values already span roughly one order of magnitude between *p*_1_ and *p*_99_), and 5 use *f* = log(1 + *x*) to compress heavy-tailed coverage distributions: observed H3K27ac, observed H3K4me3, RNA-seq + strand, RNA-seq − strand, and the super-enhancer aggregate score. The choice of *f* is made per track from the empirical value distribution observed during offline fitting (below) and fixed in the track configuration.

##### Adaptive per-track *v*_min_, *v*_max_ fitting

For every BigWig track, the scaling bounds are estimated offline by drawing 5,000 probe windows of 1,000 bp uniformly at random across the autosomes (chromosomes 1–22, restricted to positions where the window fits fully within the chromosome), reading all non-null finite values from those windows into a single sample, and computing percentiles *p*_1_, *p*_5_, *p*_50_, *p*_95_, *p*_99_, *p*_99.9_. We set *v*_min_ = *p*_1_ and *v*_max_ = *p*_99_. Tracks whose sampled values are ≥ 50% zero (sparse tracks such as gene-constraint BigWigs whose non-zero values only occur at gene regions) instead use *v*_max_ = *p*_99.9_ to prevent the clipping bound from compressing the non-zero dynamic range against a near-zero upper limit.

##### gnomAD single-nucleotide-variant allele frequency

gnomAD SNV allele frequency [21] is handled by a dedicated sparse module rather than the general continuous-normalization path because the raw data are point-based rather than dense over the genome. At positions carrying a gnomAD v4.1 SNV call with allele frequency AF, we set

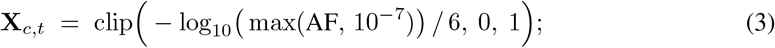

all positions not carrying a gnomAD SNV remain 0. The floor of 10^−7^ on AF prevents divergence on singleton variants; the denominator 6 corresponds to the effective upper bound of gnomAD AF resolution (log_10_(10^−6^) = 6) at current cohort size (∼10^5^). Higher signal values therefore correspond to rarer variants—the direction associated with functional constraint.

##### Categorical with graded strength

Four annotation sources attach an ordered evidence category to each interval rather than a continuous score: ClinGen gene–disease validity, mutational-constraint quantile [44], TAD boundary strength [3], and the ENCODE ChromHMM 15-state chromatin model [45]. These are rendered as single channels under a fixed ordinal map from category to scalar strength in [0, 1]. For example, `clinGenGeneDisease` assigns Definitive/Strong → 1.0, Moderate → 0.8, Limited → 0.6, down to No-known → 0.0; ChromHMM assigns Quiescent → 0 and Active TSS → 1, with 13 intermediate states on a 14-step ordinal scale; TAD boundary strength uses published stability quantiles. Where multiple annotations cover the same base pair—for example, overlapping gene isoforms with differing ClinGen validity labels—the larger strength value is retained, matching the max-reduction rule used for binary tracks. Complete category-to-strength mappings will be provided in the released codebase.

##### Variant-identity channels

Two channels are derived directly from the variant record rather than from external annotation files, and are not subject to any of the three regimes above. The *variant mask* (channel 0) takes value 0.5 inside the CNV interval for deletions and 1.0 for duplications, with value 0.0 in the 500 bp flanking padding on each side—jointly encoding variant type and variant/flank boundary. The *length channel* (channel 1) holds a log-normalized CNV length (log svlen − log 50)/(log 10^6^ log 50), clipped to [0, 1] and filled across the variant span (0 in padding); the effective length range is therefore 50 bp to 1 Mb. Both channels are programmatically injected by the tensor builder and are excluded from all percentile fitting and augmentation-group-dropout procedures.

### A.2 Model architecture

Full implementation details, layer specifications, and training configurations will be provided in the released codebase.

#### Variant-type conditioning

Across all configurations the variant type *t* ∈ DEL, DUP is injected into the window-embedding stream via a learned additive embedding applied before the sequence backbone. A lookup table **E** ∈ ℝ^2×*d*^, with rows **E**[DEL] and **E**[DUP] initialized from 𝒩 (0,, 0.02^2^), is indexed once per variant to obtain a single *d*-dimensional type vector **e**_*t*_ = **E**[*t*]. This vector is broadcast across all *K* windows and added to each window embedding produced by the encoder,

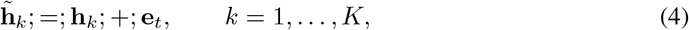

so that the subsequent sequence backbone and variant aggregator operate on type-informed representations. The embedding table is trained jointly with the rest of the model and adds only 2*d* parameters.

#### Classification head

The aggregated variant representation is mapped to a pathogenicity logit through a two-layer MLP with ReLU activations and dropout.

##### A.2.1 Self-supervised VAE window encoder

###### Overview

The self-supervised encoder used by Tesseract-VAE is a convolutional variational autoencoder with a cross-attention bottleneck, pretrained on unlabeled 500 bp windows under a masked-reconstruction objective. After pretraining, the encoder is frozen and used as a fixed feature extractor for downstream classification.

###### Input representation

Each 500 bp window is presented to the pretraining model under the channelas-height convention: the 78 × 500 annotation matrix is reshaped to a (1, 78, 500) single-channel image so that the 78 annotation tracks act as the spatial height axis and 500 bp of genomic position form the width axis. Two-dimensional convolution kernels are therefore free to mix across tracks as well as along position, matching the encoder used in the end-to-end pipeline (Methods 3.2).

###### Architecture

The pretraining architecture consists of four components: (i) a CNN tokenizer shared with the end-to-end encoder that converts the input image into a spatial feature sequence; (ii) a multi-head cross-attention bottleneck [29] that compresses the spatial sequence into *K* = 64 latent tokens; (iii) variational posterior heads that parameterize a Gaussian latent distribution and apply standard reparameterization; and (iv) a lightweight convolutional decoder that reconstructs the masked input image from the latent representation. Full architectural details will be provided in the released codebase.

###### Masked-reconstruction objective

Before tokenization the input is randomly masked on a fixed patch grid, in the spirit of masked-autoencoder pretraining [28]. The masking grid has patch size 6 × 50 (track × position), yielding 13 × 10 = 130 patches per window; a fraction *r* = 0.5 of these patches is drawn uniformly without replacement and set to zero per example. Masking is active only when the model is in training mode; evaluation and feature-extraction forward passes operate on the full unmasked input.

The per-example loss is

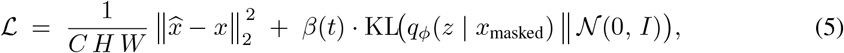

where the reconstruction term is computed against the full unmasked target image, and the KL divergence term is annealed linearly during early training.

###### Pretraining corpus

Windows are extracted exclusively from the training and validation chromosomes of the chromosome-stratified PhenoSV partition, excluding all held-out test and independent evaluation cohorts by construction. Tiling variants into non-overlapping 500 bp windows yields a pretraining corpus of approximately 4.1 million windows, substantially larger than the labeled CNV corpus and motivating the self-supervised design.

###### Optimization

Pretraining uses AdamW [34] with cosine learning-rate decay, mixed-precision training, gradient clipping, and early stopping on validation reconstruction loss. Full optimization hyperparameters will be provided in the released codebase.

###### Feature extraction and freezing

After pretraining, the decoder and variance head are discarded and the remaining encoder components are frozen for all downstream experiments. Each 500 bp window is embedded independently into 64 latent tokens using the posterior mean. For downstream classification, these tokens are mean-pooled into a single window embedding **h**_*k*_ ∈ ℝ^128^ before sequence modeling.

### A.3 Data sources and preprocessing

#### Coding/noncoding labels

Following Xu et al. [18], a variant is labeled *coding* if it overlaps at least one base pair of a protein-coding exon in the GENCODE v48 annotation [46] and *noncoding* otherwise. We re-derive this label from a single GENCODE release so that coding/noncoding partitions are defined identically for TESSERACT-CNV and every baseline.

##### A.3.1 Training data and held-out test

The training and testing data follow the CNV corpus construction and chromosome-based split protocol of PhenoSV [18]. The corpus is derived from ClinVar full release 02/2022 [32], DECIPHER v11.15 [33], SvAnna [12], and NCBI common structural-variant calls [47]. Variants are filtered by clinical significance, SV type, size, genome build, reciprocal-overlap redundancy, and label conflicts following the PhenoSV preprocessing protocol. For reproducibility, we report the source datasets, split definitions, and preprocessing protocol; third-party clinical-genomics records can be obtained from their original providers under the corresponding usage terms.

##### A.3.2 Independent test preprocessing

Starting from the DECIPHER v11.15 release [33] we apply the same label-consistency and size filtering used by PhenoSV [18], followed by the two additional leakage-control steps summarized in the main text (Section 3.3). The end-to-end pipeline is:

1. **SV-type restriction**. We retain only deletion and duplication events.
2. **Length filter**. We retain variants longer than or equal to 50 bp.
3. **Clinical-label harmonization**. We retain only variants labeled as *Pathogenic, Likely pathogenic, Benign*, or *Likely benign*, discarding variants with uncertain, conflicting, or missing clinical significance annotations. For binary evaluation, pathogenic and likely pathogenic labels are merged into the positive class, and benign and likely benign labels into the negative class.
4. **Reciprocal-overlap leakage filtering**. We remove DECIPHER variants with ≥ 90% reciprocal overlap against same-label training variants or ≥ 85% reciprocal overlap against cross-label training variants.

The resulting coding evaluation cohort contains 3,041 variants and serves as the primary independent benchmark reported in Section 4. Its breakdown by SV type and label is summarized in Table 3.

##### A.3.3 Per-split CNV counts

### A.4 Training and evaluation protocol

#### A.4.1 Training hyperparameters

##### Supervised classifier training

Supervised training uses AdamW [34] with learning rate 1 × 10^−4^, weight decay 0.05, and (*β*_1_, *β*_2_) = (0.9, 0.95). We use cosine learning-rate decay with 6.25% linear warm-up, mixed-precision (FP16) training, and early stopping on validation AUROC with patience 15 for a maximum of 60 epochs. Training is performed on 4 NVIDIA A10G GPUs.

##### Input-side regularization

For Tesseract-E2E, we apply annotation-group dropout with probability 0.15 independently across the 14 annotation groups. For Tesseract-VAE, we apply dropout with rate 0.1 to the window embeddings before sequence modeling.

##### Self-supervised pretraining (VAE)

The Tesseract-VAE encoder is pretrained separately under the masked-reconstruction objective described in Appendix A.2.1. Pretraining uses AdamW with (*β*_1_, *β*_2_) = (0.9, 0.95), FP16 training, and early stopping on validation reconstruction loss with patience 15 for a maximum of 60 epochs. Downstream supervised training consumes only the frozen pretrained encoder through precomputed latent embeddings and otherwise follows the same optimization protocol described above.

##### Compute resources

Training was performed on NVIDIA A10G GPUs with mixed-precision training. The self-supervised MCAB-VAE pretraining run used a single A10G GPU and required approximately 6.0 wall-clock hours, stopping at epoch 22 once validation reconstruction loss had plateaued. The supervised Tesseract-E2E headline model was trained with distributed data parallelism on 4 A10G GPUs with an effective batch size of 32 and converged in approximately 2.6 wall-clock hours, corresponding to approximately 10.4 GPU-hours, with 27 epochs completed before early stopping on validation AUROC. The downstream Tesseract-VAE classifier, trained on cached frozen VAE latents, required approximately 15 minutes on a single A10G GPU, corresponding to approximately 0.25 GPU-hours, with 32 epochs completed before early stopping. Including the one-time VAE pretraining cost, the Tesseract-VAE training path required approximately 6.3 GPU-hours for the headline configuration with batch size of 64. VAE latents were stored on a local storage and reused for downstream supervised training, ablations, and segmented inference.

#### A.4.2 Baseline eligibility

The benchmark comparison in Section 4.2.1 is constrained by what each published baseline saw during training and by what each tool can score at evaluation time. We therefore report every baseline only on evaluation splits that are outside its training set and supported by its input requirements.

Unless otherwise stated, each baseline was run using the default inference settings and pretrained models provided by the corresponding authors’ code repositories or documentation. We did not tune baseline hyperparameters on our evaluation splits. For probabilistic binary comparisons, we use each tool’s reported pathogenicity score where available; for CNVoyant[16], which outputs a three-class distribution over {`BENIGN, VUS, PATHOGENIC`}, we use *P* (`PATHOGENIC`) for ranking metrics and its native three-class argmax for native operating-point reporting.

##### CNVoyant

CNVoyant [16] is trained on ClinVar coding CNVs under five-fold cross-validation. We therefore do not report it on the ClinVar coding split used for the in-distribution benchmark, because that split is drawn from the same source domain and would not constitute a clean held-out comparison to its published training setup. We do report CNVoyant on DECIPHER IndT, which is the primary external benchmark, and on the ClinVar noncoding split, whose pathogenic variants are curated across ClinVar, DECIPHER, and SvAnna with benign counterparts from NCBI common-SV calls rather than from the ClinVar coding pool used for CNVoyant training.

##### TADA

TADA [15] is trained on DECIPHER-derived data and is therefore evaluated only on the ClinVar test splits in our comparison. On the noncoding ClinVar subset, TADA is additionally limited by its TAD-based feature coverage, so the reported numbers use the subset of variants for which the tool returns a score.

##### StrVCTVRE

StrVCTVRE [17] is trained on ClinVar and is therefore evaluated only on the external DECIPHER IndT cohort. Its reported sample size is smaller than the full cohort because some variants fall outside the tool’s scoring coverage.

##### PhenoSV

PhenoSV [18] is trained on the same chromosome-stratified corpus used to train Tesseract. We report it on every evaluation split because it is the most directly comparable prior method, but its numbers on the held-out PhenoSV-derived test split should be interpreted as an in-distribution reference rather than an external comparison.

### A.5 Benchmarking details

#### Conventions

All benchmarking tables in this subsection report: AUROC and average precision (AP) as ranking metrics; *F*_1_, precision, and recall as pathogenic-class operating-point metrics at the default decision threshold (*P* (pathogenic) ≥ 0.5, or argmax for multiclass classifiers; see §4.1); Brier score computed on *P* (pathogenic); and expected calibration error (ECE) with 15 equal-width probability bins. Tesseract configurations are reported under both inference regimes (segmented at *S* = 120 and full-length); published baselines are single-configuration. †marks an in-distribution reference (PhenoSV shares the training corpus of Tesseract). ‡ marks a scoring-convention caveat explained in the relevant table.

**Table 4:** Panel A (Figure 3A): Held-out test coding, *n* = 2,559.

| Model (regime) | AUROC | AP | F<sub>1</sub> | Precision | Recall | Brier | ECE |
| --- | --- | --- | --- | --- | --- | --- | --- |
| TESSERACT-E2E, $S = 120$ | 0.942 | 0.947 | 0.870 | 0.795 | **0.960** | 0.118 | 0.118 |
| TESSERACT-E2E, full-length | 0.954 | 0.953 | **0.900** | 0.866 | 0.936 | 0.086 | 0.060 |
| TESSERACT-VAE, $S = 120$ | **0.956** | **0.965** | **0.900** | **0.894** | 0.907 | **0.083** | **0.053** |
| TESSERACT-VAE, full-length | 0.942 | 0.955 | 0.866 | 0.817 | 0.921 | 0.115 | 0.093 |
| PhenoSV<sup>†</sup> | 0.853 | 0.869 | 0.802 | 0.722 | 0.901 | 0.172 | 0.109 |
| TADA | 0.694 | 0.734 | 0.719 | 0.595 | 0.908 | 0.233 | 0.120 |

**Table 5:** Panel B (Figure 3B): Held-out test noncoding, *n* = 114. TADA is scored on *n* = 105 variants due to TAD-boundary coverage. ^‡^CNVoyant assigns BENIGN to every noncoding variant under its native three-class argmax classifier, so F_1_/precision/recall collapse to 0.000. Ranking metrics (AUROC, AP) remain strong because *P* (pathogenic) still orders pathogenic above benign; the collapse reflects a calibration gap between CNVoyant’s coding-trained output head and the noncoding distribution, not a scoring-convention mismatch.

| Model (regime) | AUROC | AP | F<sub>1</sub> | Precision | Recall | Brier | ECE |
| --- | --- | --- | --- | --- | --- | --- | --- |
| TESSERACT-E2E, $S = 120$ | **0.936** | **0.950** | 0.827 | 0.915 | **0.754** | **0.109** | **0.116** |
| TESSERACT-E2E, full-length | 0.926 | 0.942 | **0.835** | 0.935 | **0.754** | 0.129 | 0.153 |
| TESSERACT-VAE, $S = 120$ | 0.930 | 0.943 | 0.659 | **1.000** | 0.491 | 0.198 | 0.232 |
| TESSERACT-VAE, full-length | 0.885 | 0.892 | 0.667 | 0.909 | 0.526 | 0.190 | 0.200 |
| CNVoyant<sup>‡</sup> | 0.885 | 0.927 | 0.000 | 0.000 | 0.000 | 0.396 | 0.442 |
| PhenoSV<sup>†</sup> | 0.862 | 0.865 | 0.769 | 0.851 | 0.702 | 0.176 | 0.138 |
| TADA ($n = 105$) | 0.405 | 0.582 | 0.235 | 0.727 | 0.140 | 0.358 | 0.406 |

**Table 6:** CNVoyant score distribution on the held-out noncoding benchmark. BEN = benign, PATH = pathogenic, and VUS = variant of uncertain significance. The pathogenic-probability head never becomes the dominant class: AUROC remains informative because *P* (BEN) is lower on pathogenic variants, but the native three-class argmax assigns every variant to BEN.

| True label | $n$ | Mean $P(\text{BEN})$ | Mean $P(\text{VUS})$ | Mean $P(\text{PATH})$ | Max $P(\text{PATH})$ | Argmax BEN/VUS/PATH |
| --- | --- | --- | --- | --- | --- | --- |
| Benign | 57 | 0.98 | 0.02 | 0.01 | 0.02 | 57/0/0 |
| Pathogenic | 59 | 0.77 | 0.10 | 0.12 | 0.43 | 59/0/0 |

**Table 7:** Panel C (Figure 3C): Independent test, *n* = 3,041. StrVCTVRE is scored on *n* = 2,964 variants due to its own coverage constraints. CNVoyant is scored on *P* (pathogenic) for AUROC, AP, Brier, and ECE and on its native argmax for *F*_1_, Precision, and Recall.

| Model (regime) | AUROC | AP | $F_1$ | Precision | Recall | Brier | ECE |
| --- | --- | --- | --- | --- | --- | --- | --- |
| TESSERACT-E2E, $S = 120$ | 0.826 | 0.901 | 0.778 | 0.852 | 0.716 | 0.199 | 0.160 |
| TESSERACT-E2E, full-length | 0.808 | 0.893 | 0.724 | 0.894 | 0.608 | 0.248 | 0.227 |
| TESSERACT-VAE, $S = 120$ | **0.835** | 0.899 | 0.752 | 0.880 | 0.656 | 0.214 | 0.190 |
| TESSERACT-VAE, full-length | 0.815 | 0.883 | **0.809** | 0.800 | **0.817** | **0.185** | 0.133 |
| CNVoyant | 0.829 | **0.906** | 0.468 | **0.981** | 0.307 | 0.376 | 0.426 |
| PhenoSV | 0.744 | 0.826 | 0.782 | 0.770 | 0.795 | 0.210 | 0.118 |
| StrVCTVRE ($n = 2,964$) | 0.758 | 0.862 | 0.759 | 0.784 | 0.735 | 0.197 | **0.100** |

### A.6 Architecture ablation details

**Table 8:** Panel A (Figure 4A): architecture-ablation grid on independent test, *n* = 3,041. Full 2 × 3 × 3 grid of window-encoder regime × sequence backbone × pooling operator, reporting each configuration under segmented inference (*S* = 120) and full-length inference on consecutive rows. Regime: end-to-end (E2E) 2D CNN vs frozen pretrained VAE encoder. Pooling: attention-based MIL (ABMIL), masked mean, max. Metrics and conventions are as in Table 7. The two headline configurations of the main text are the best cell on each side of the encoder-regime axis at *S* = 120: Tesseract-E2E = E2E + no backbone + ABMIL; Tesseract-VAE = VAE + Transformer-4L + ABMIL.

| Regime | Backbone | Pool | Inference | AUROC | AP | F<sub>1</sub> | P | R | Brier | ECE |
| --- | --- | --- | --- | --- | --- | --- | --- | --- | --- | --- |
| E2E | None | ABMIL | $S = 120$ | 0.826 | **0.901** | 0.778 | 0.852 | 0.716 | 0.199 | 0.160 |
| E2E | None | ABMIL | full | 0.808 | 0.893 | 0.724 | 0.894 | 0.608 | 0.248 | 0.227 |
| E2E | None | mean | $S = 120$ | 0.823 | 0.896 | 0.757 | 0.868 | 0.671 | 0.211 | 0.181 |
| E2E | None | mean | full | 0.808 | 0.890 | 0.676 | 0.905 | 0.539 | 0.270 | 0.264 |
| E2E | None | max | $S = 120$ | 0.805 | 0.882 | 0.787 | 0.811 | 0.764 | 0.205 | 0.161 |
| E2E | None | max | full | 0.822 | 0.896 | 0.713 | 0.905 | 0.589 | 0.253 | 0.240 |
| E2E | Trans-4L | ABMIL | $S = 120$ | 0.821 | 0.896 | 0.790 | 0.843 | 0.743 | 0.192 | 0.139 |
| E2E | Trans-4L | ABMIL | full | 0.792 | 0.881 | 0.695 | 0.895 | 0.568 | 0.265 | 0.250 |
| E2E | Trans-4L | mean | $S = 120$ | 0.829 | 0.897 | 0.800 | 0.836 | 0.767 | 0.210 | 0.191 |
| E2E | Trans-4L | mean | full | 0.811 | 0.889 | 0.707 | 0.912 | 0.577 | 0.276 | 0.268 |
| E2E | Trans-4L | max | $S = 120$ | 0.816 | 0.893 | 0.777 | 0.846 | 0.719 | 0.219 | 0.184 |
| E2E | Trans-4L | max | full | 0.809 | 0.890 | 0.710 | 0.907 | 0.583 | 0.273 | 0.263 |
| E2E | Mamba-4L | ABMIL | $S = 120$ | 0.822 | 0.899 | 0.739 | 0.861 | 0.648 | 0.231 | 0.204 |
| E2E | Mamba-4L | ABMIL | full | 0.810 | 0.892 | 0.655 | **0.927** | 0.507 | 0.301 | 0.298 |
| E2E | Mamba-4L | mean | $S = 120$ | 0.817 | 0.889 | 0.753 | 0.866 | 0.667 | 0.207 | 0.177 |
| E2E | Mamba-4L | mean | full | 0.803 | 0.879 | 0.665 | 0.907 | 0.525 | 0.270 | 0.264 |
| E2E | Mamba-4L | max | $S = 120$ | 0.811 | 0.882 | 0.712 | 0.889 | 0.594 | 0.256 | 0.245 |
| E2E | Mamba-4L | max | full | 0.819 | 0.894 | 0.685 | 0.916 | 0.547 | 0.282 | 0.279 |
| VAE | None | ABMIL | $S = 120$ | 0.806 | 0.878 | 0.742 | 0.868 | 0.648 | 0.210 | 0.161 |
| VAE | None | ABMIL | full | 0.788 | 0.871 | 0.635 | 0.904 | 0.490 | 0.268 | 0.269 |
| VAE | None | mean | $S = 120$ | 0.795 | 0.869 | 0.723 | 0.861 | 0.623 | 0.220 | 0.175 |
| VAE | None | mean | full | 0.774 | 0.857 | 0.602 | 0.896 | 0.454 | 0.281 | 0.283 |
| VAE | None | max | $S = 120$ | 0.798 | 0.870 | 0.753 | 0.846 | 0.678 | 0.199 | 0.136 |
| VAE | None | max | full | 0.756 | 0.850 | 0.581 | 0.885 | 0.433 | 0.290 | 0.292 |
| VAE | Trans-4L | ABMIL | $S = 120$ | **0.835** | 0.899 | 0.752 | 0.880 | 0.656 | 0.214 | 0.190 |
| VAE | Trans-4L | ABMIL | full | 0.815 | 0.883 | **0.809** | 0.800 | **0.817** | **0.185** | **0.133** |
| VAE | Trans-4L | mean | $S = 120$ | 0.822 | 0.884 | 0.783 | 0.839 | 0.735 | 0.190 | 0.139 |
| VAE | Trans-4L | mean | full | 0.787 | 0.851 | 0.793 | 0.806 | 0.781 | 0.200 | 0.158 |
| VAE | Trans-4L | max | $S = 120$ | 0.831 | 0.899 | 0.731 | 0.893 | 0.619 | 0.221 | 0.198 |
| VAE | Trans-4L | max | full | 0.802 | 0.867 | 0.806 | 0.802 | 0.809 | 0.197 | 0.165 |
| VAE | Mamba-4L | ABMIL | $S = 120$ | 0.816 | 0.891 | 0.770 | 0.848 | 0.705 | 0.209 | 0.167 |
| VAE | Mamba-4L | ABMIL | full | 0.802 | 0.881 | 0.689 | 0.899 | 0.558 | 0.265 | 0.246 |
| VAE | Mamba-4L | mean | $S = 120$ | 0.822 | 0.889 | 0.743 | 0.877 | 0.644 | 0.225 | 0.197 |
| VAE | Mamba-4L | mean | full | 0.809 | 0.879 | 0.628 | 0.911 | 0.479 | 0.302 | 0.309 |
| VAE | Mamba-4L | max | $S = 120$ | 0.821 | 0.887 | 0.754 | 0.866 | 0.667 | 0.210 | 0.173 |
| VAE | Mamba-4L | max | full | 0.803 | 0.882 | 0.666 | 0.909 | 0.525 | 0.274 | 0.267 |

**Table 9:**
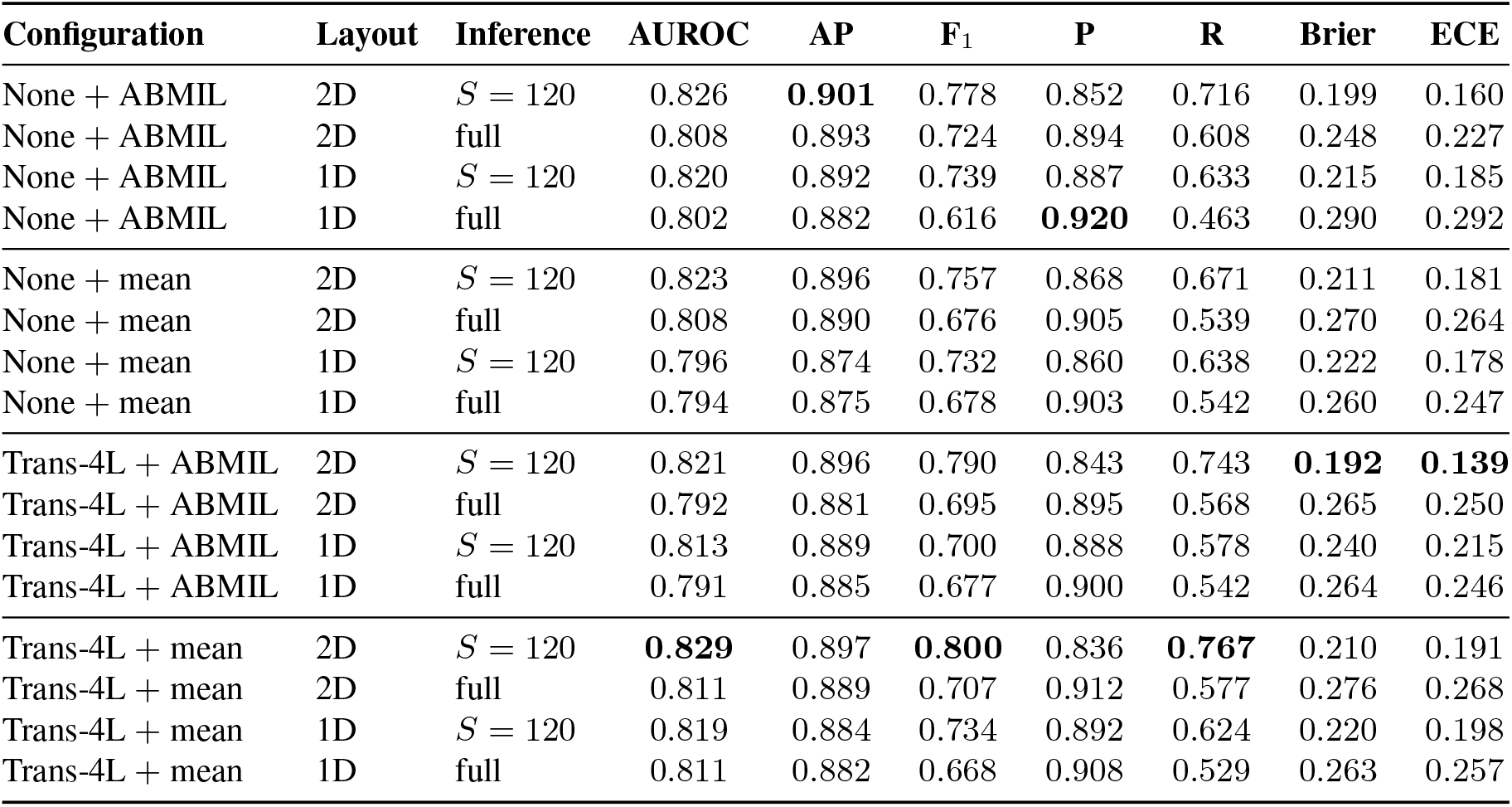
Panel B (Figure 4B): Tesseract-E2E 2D channel-as-height versus 1D per-channel encoder on independent test, *n* = 3,041. The four strongest Panel A configurations are paired against their 1D counterparts; each (configuration, layout) pair is reported under segmented inference (*S* = 120) and full-length inference on consecutive rows. Metrics and conventions are as in Table 7.

| Configuration | Layout | Inference | AUROC | AP | F<sub>1</sub> | P | R | Brier | ECE |
| --- | --- | --- | --- | --- | --- | --- | --- | --- | --- |
| None + ABMIL | 2D | $S = 120$ | 0.826 | **0.901** | 0.778 | 0.852 | 0.716 | 0.199 | 0.160 |
| None + ABMIL | 2D | full | 0.808 | 0.893 | 0.724 | 0.894 | 0.608 | 0.248 | 0.227 |
| None + ABMIL | 1D | $S = 120$ | 0.820 | 0.892 | 0.739 | 0.887 | 0.633 | 0.215 | 0.185 |
| None + ABMIL | 1D | full | 0.802 | 0.882 | 0.616 | **0.920** | 0.463 | 0.290 | 0.292 |
| None + mean | 2D | $S = 120$ | 0.823 | 0.896 | 0.757 | 0.868 | 0.671 | 0.211 | 0.181 |
| None + mean | 2D | full | 0.808 | 0.890 | 0.676 | 0.905 | 0.539 | 0.270 | 0.264 |
| None + mean | 1D | $S = 120$ | 0.796 | 0.874 | 0.732 | 0.860 | 0.638 | 0.222 | 0.178 |
| None + mean | 1D | full | 0.794 | 0.875 | 0.678 | 0.903 | 0.542 | 0.260 | 0.247 |
| Trans-4L + ABMIL | 2D | $S = 120$ | 0.821 | 0.896 | 0.790 | 0.843 | 0.743 | **0.192** | **0.139** |
| Trans-4L + ABMIL | 2D | full | 0.792 | 0.881 | 0.695 | 0.895 | 0.568 | 0.265 | 0.250 |
| Trans-4L + ABMIL | 1D | $S = 120$ | 0.813 | 0.889 | 0.700 | 0.888 | 0.578 | 0.240 | 0.215 |
| Trans-4L + ABMIL | 1D | full | 0.791 | 0.885 | 0.677 | 0.900 | 0.542 | 0.264 | 0.246 |
| Trans-4L + mean | 2D | $S = 120$ | **0.829** | 0.897 | **0.800** | 0.836 | **0.767** | 0.210 | 0.191 |
| Trans-4L + mean | 2D | full | 0.811 | 0.889 | 0.707 | 0.912 | 0.577 | 0.276 | 0.268 |
| Trans-4L + mean | 1D | $S = 120$ | 0.819 | 0.884 | 0.734 | 0.892 | 0.624 | 0.220 | 0.198 |
| Trans-4L + mean | 1D | full | 0.811 | 0.882 | 0.668 | 0.908 | 0.529 | 0.263 | 0.257 |

### A.7 Training-window count sweep

**Table 10:** Training-window-count sweep on independent test, *n* = 3,041. We retrain the Tesseract-VAE configuration (VAE + Transformer-4L + ABMIL) at five training-time window counts *K* ∈ {80, 120, 200, 400, 1000} on the same training split with every other hyperparameter fixed. Each model is evaluated at its training-matched segmented-inference regime (except *K* = 1000, which is evaluated at *S* = 800 as the nearest available segment size in the sweep) and at full-length inference. Metrics and conventions are as in Table 7. The *K* = 120 configuration is the headline Tesseract-VAE.

| Training $K$ | Inference | AUROC | AP | $F_1$ | P | R | Brier | ECE |
| --- | --- | --- | --- | --- | --- | --- | --- | --- |
| $K = 80$ | $S = 80$ (matched) | **0.836** | **0.901** | 0.786 | 0.867 | 0.720 | 0.190 | 0.149 |
| $K = 80$ | full | 0.807 | 0.873 | 0.810 | 0.799 | 0.822 | **0.185** | 0.134 |
| $K = 120$ | $S = 120$ (matched) | 0.835 | 0.899 | 0.752 | 0.880 | 0.656 | 0.214 | 0.190 |
| $K = 120$ | full | 0.815 | 0.883 | 0.809 | 0.800 | 0.817 | **0.185** | **0.133** |
| $K = 200$ | $S = 200$ (matched) | 0.833 | 0.900 | 0.752 | 0.891 | 0.650 | 0.206 | 0.175 |
| $K = 200$ | full | 0.807 | 0.876 | **0.825** | 0.784 | **0.871** | 0.186 | 0.135 |
| $K = 400$ | $S = 400$ (matched) | 0.829 | 0.898 | 0.714 | 0.900 | 0.592 | 0.244 | 0.231 |
| $K = 400$ | full | 0.801 | 0.868 | 0.767 | 0.834 | 0.711 | 0.230 | 0.215 |
| $K = 1000$ | $S = 800$ (nearest) | 0.819 | 0.891 | 0.679 | **0.909** | 0.542 | 0.267 | 0.262 |
| $K = 1000$ | full | 0.817 | 0.887 | 0.720 | 0.878 | 0.610 | 0.248 | 0.233 |

### A.8 Full metric tables for Figure 5

**Table 11:** Operating-point and ranking metrics by model × SV type × segment size on Independent test. All numbers computed at decision threshold 0.5 with `max` aggregation over per-segment probabilities. Rows grouped by segment size *S*; within each *S*-block, DEL and DUP panels report metrics for both Tesseract-E2E and Tesseract-VAE, and an **ALL** row reports metrics over the full coding cohort (*n* = 3,041; DEL: *n* = 1,646, DUP: *n* = 1,395). Bolded value per column indicates the per-model maximum across the segment-size sweep for that SV group. The training-matched *S* = 120 is the overall AUROC and AP optimum for Tesseract-VAE (0.835, 0.899), while fine *S* shifts the operating point toward recall (Tesseract-VAE F_1_/R peak at *S* = 20: 0.822/0.917). On DEL the two models track each other within 0.05 on every metric at every *S*. On DUP the story diverges at large *S*: at *S* = 1200 and at full-length, Tesseract-VAE recovers to a balanced DUP operating point (F_1_ ≈ 0.68, R≈ 0.69), whereas Tesseract-E2E collapses on recall (F_1_ ≈ 0.30, R≈ 0.19). Numerical backing for Figure 5 and Section 4.3.1.

| $S$ | SV | AUROC | | AP | | $F_1$ | | Precision | | Recall | |
| --- | --- | --- | --- | --- | --- | --- | --- | --- | --- | --- | --- |
|  |  | E2E | VAE | E2E | VAE | E2E | VAE | E2E | VAE | E2E | VAE |
| $S = 10$ | DEL | 0.844 | 0.800 | 0.949 | 0.920 | 0.893 | 0.888 | 0.831 | 0.851 | **0.966** | 0.928 |
|  | DUP | 0.726 | 0.728 | 0.705 | 0.680 | **0.679** | 0.711 | 0.641 | 0.598 | **0.722** | 0.878 |
|  | ALL | 0.819 | 0.778 | 0.896 | 0.848 | **0.819** | 0.819 | 0.766 | 0.745 | **0.880** | 0.910 |
| $S = 20$ | DEL | 0.849 | 0.799 | **0.951** | 0.918 | **0.897** | **0.892** | 0.847 | 0.856 | 0.953 | **0.932** |
|  | DUP | 0.730 | 0.724 | 0.712 | 0.680 | 0.641 | **0.713** | 0.668 | 0.595 | 0.617 | **0.889** |
|  | ALL | 0.823 | 0.781 | 0.899 | 0.852 | 0.813 | **0.822** | 0.792 | 0.745 | 0.836 | **0.917** |
| $S = 40$ | DEL | 0.849 | 0.822 | **0.951** | 0.931 | 0.894 | 0.877 | 0.861 | 0.868 | 0.930 | 0.887 |
|  | DUP | 0.746 | 0.747 | 0.729 | 0.703 | 0.608 | 0.696 | 0.718 | 0.664 | 0.528 | 0.732 |
|  | ALL | 0.828 | 0.810 | 0.901 | 0.876 | 0.806 | 0.813 | 0.823 | 0.794 | 0.790 | 0.833 |
| $S = 80$ | DEL | **0.851** | 0.831 | **0.951** | 0.936 | 0.888 | 0.875 | 0.868 | 0.878 | 0.909 | 0.873 |
|  | DUP | **0.749** | 0.761 | **0.739** | 0.727 | 0.555 | 0.635 | 0.743 | 0.700 | 0.443 | 0.582 |
|  | ALL | **0.830** | 0.825 | **0.903** | 0.890 | 0.790 | 0.796 | 0.839 | 0.823 | 0.746 | 0.772 |
| $S = 120$ | DEL | 0.850 | **0.846** | **0.951** | **0.942** | 0.885 | 0.853 | 0.877 | **0.908** | 0.892 | 0.805 |
|  | DUP | 0.741 | 0.764 | 0.729 | **0.747** | 0.512 | 0.510 | 0.759 | 0.785 | 0.386 | 0.378 |
|  | ALL | 0.826 | **0.835** | 0.901 | **0.899** | 0.778 | 0.752 | 0.852 | **0.880** | 0.716 | 0.656 |
| $S = 200$ | DEL | **0.851** | 0.839 | **0.951** | **0.942** | 0.882 | 0.867 | 0.887 | 0.895 | 0.877 | 0.840 |
|  | DUP | 0.744 | 0.763 | 0.738 | 0.743 | 0.482 | 0.504 | 0.832 | **0.791** | 0.340 | 0.370 |
|  | ALL | 0.827 | 0.830 | 0.902 | 0.897 | 0.772 | 0.762 | 0.877 | 0.873 | 0.689 | 0.676 |
| $S = 400$ | DEL | 0.846 | 0.822 | 0.949 | 0.935 | 0.875 | 0.877 | 0.894 | 0.870 | 0.856 | 0.884 |
|  | DUP | 0.729 | **0.771** | 0.725 | 0.740 | 0.401 | 0.568 | 0.838 | 0.751 | 0.264 | 0.456 |
|  | ALL | 0.820 | 0.825 | 0.898 | 0.891 | 0.749 | 0.784 | 0.886 | 0.841 | 0.650 | 0.735 |
| $S = 800$ | DEL | 0.842 | 0.815 | 0.948 | 0.933 | 0.869 | 0.876 | **0.897** | 0.866 | 0.841 | 0.887 |
|  | DUP | 0.713 | 0.763 | 0.711 | 0.730 | 0.333 | 0.683 | 0.845 | 0.676 | 0.207 | 0.691 |
|  | ALL | 0.811 | 0.816 | 0.894 | 0.885 | 0.731 | 0.809 | 0.891 | 0.800 | 0.620 | 0.818 |
| $S = 1200$ | DEL | 0.841 | 0.815 | 0.947 | 0.932 | 0.866 | 0.876 | **0.897** | 0.866 | 0.838 | 0.887 |
|  | DUP | 0.708 | 0.763 | 0.706 | 0.729 | 0.310 | 0.682 | 0.855 | 0.677 | 0.190 | 0.687 |
|  | ALL | 0.809 | 0.817 | 0.893 | 0.885 | 0.726 | 0.809 | 0.892 | 0.801 | 0.611 | 0.817 |
| full | DEL | 0.841 | 0.813 | 0.948 | 0.931 | 0.865 | 0.876 | **0.897** | 0.866 | 0.835 | 0.887 |
|  | DUP | 0.707 | 0.763 | 0.709 | 0.725 | 0.304 | 0.682 | **0.875** | 0.676 | 0.184 | 0.688 |
|  | ALL | 0.808 | 0.815 | 0.893 | 0.883 | 0.724 | 0.809 | **0.894** | 0.800 | 0.608 | 0.817 |

**Table 12:** Aggregator sweep at representative segment sizes, Tesseract-VAE, Independent test. Per-segment probabilities are collapsed to a variant-level score via one of six aggregators: `max` or top-*k* mean for *k* ∈ {2, 3, 5, 10, 20}. Three segment sizes are shown: two fine (*S* = 10, *S* = 20) where aggregator choice has its largest effect, and the training-matched *S* = 120 where aggregator choice is negligible. Best value per metric within each *S*-block is in bold. At *S* = 120 the AUROC and AP spread across aggregators is ≤ 0.006, confirming that aggregator choice is second-order at the training-matched regime. At fine *S*, `max` wins F_1_ and recall by wide margins while top-*k* mean wins AUROC, AP, and precision; the aggregator therefore acts as a second-order precision–recall control at fixed *S*.

| $S$ | Aggregator | AUROC | AP | $F_1$ | Precision | Recall |
| --- | --- | --- | --- | --- | --- | --- |
| $S = 10$ | max | 0.778 | 0.848 | **0.819** | 0.745 | **0.910** |
|  | top-2 mean | 0.784 | 0.854 | 0.818 | 0.756 | 0.891 |
|  | top-3 mean | 0.786 | 0.856 | 0.815 | 0.766 | 0.870 |
|  | top-5 mean | 0.790 | 0.859 | 0.809 | 0.781 | 0.840 |
|  | top-10 mean | 0.795 | 0.863 | 0.796 | 0.801 | 0.791 |
|  | top-20 mean | **0.802** | **0.870** | 0.764 | **0.829** | 0.708 |
| $S = 20$ | max | 0.781 | 0.852 | **0.822** | 0.745 | **0.917** |
|  | top-2 mean | 0.787 | 0.859 | 0.820 | 0.754 | 0.899 |
|  | top-3 mean | 0.789 | 0.860 | 0.816 | 0.762 | 0.878 |
|  | top-5 mean | 0.791 | 0.862 | 0.812 | 0.776 | 0.852 |
|  | top-10 mean | 0.797 | 0.867 | 0.800 | 0.801 | 0.799 |
|  | top-20 mean | **0.798** | **0.869** | 0.770 | **0.826** | 0.722 |
| $S = 120$ | max | 0.835 | 0.899 | **0.752** | 0.880 | **0.656** |
|  | top-2 mean | **0.836** | 0.900 | 0.733 | 0.895 | 0.621 |
|  | top-3 mean | **0.836** | **0.902** | 0.716 | 0.901 | 0.595 |
|  | top-5 mean | 0.834 | 0.900 | 0.689 | 0.917 | 0.552 |
|  | top-10 mean | 0.832 | 0.897 | 0.651 | **0.925** | 0.503 |
|  | top-20 mean | 0.831 | 0.896 | 0.642 | **0.925** | 0.491 |

**Table 13:**
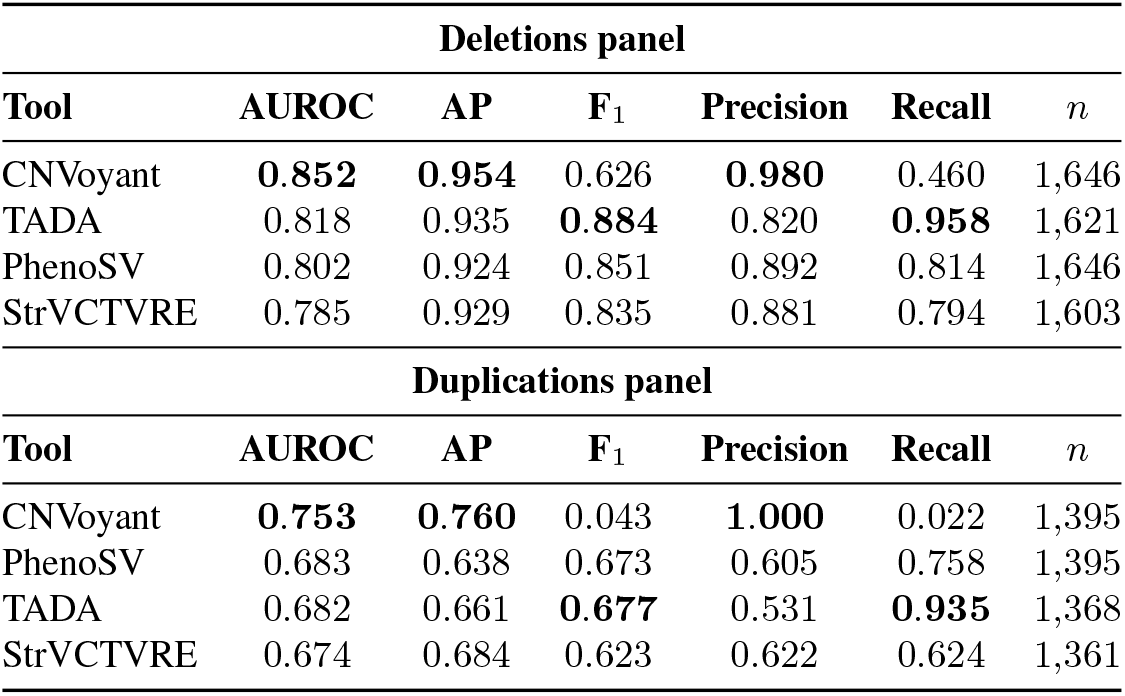
DEL/DUP split on independent test: published baselines. Numerical backing for the DEL/DUP asymmetry claim in Section 4.3.1. Each tool’s probabilistic score is thresholded at 0.5; CNVoyant’s class-probability for `PATHOGENIC` is used. Rows are sorted by AUROC within each panel. Sample sizes vary where a tool drops variants that fail its internal filters.

| Deletions panel |  |  |  |  |  |  |
| --- | --- | --- | --- | --- | --- | --- |
| Tool | AUROC | AP | F<sub>1</sub> | Precision | Recall | *n* |
| CNVoyant | **0.852** | **0.954** | 0.626 | **0.980** | 0.460 | 1,646 |
| TADA | 0.818 | 0.935 | **0.884** | 0.820 | **0.958** | 1,621 |
| PhenoSV | 0.802 | 0.924 | 0.851 | 0.892 | 0.814 | 1,646 |
| StrVCTVRE | 0.785 | 0.929 | 0.835 | 0.881 | 0.794 | 1,603 |
| Duplications panel |  |  |  |  |  |  |
| Tool | AUROC | AP | F<sub>1</sub> | Precision | Recall | *n* |
| CNVoyant | **0.753** | **0.760** | 0.043 | **1.000** | 0.022 | 1,395 |
| PhenoSV | 0.683 | 0.638 | 0.673 | 0.605 | 0.758 | 1,395 |
| TADA | 0.682 | 0.661 | **0.677** | 0.531 | **0.935** | 1,368 |
| StrVCTVRE | 0.674 | 0.684 | 0.623 | 0.622 | 0.624 | 1,361 |

### A.9 Segment-level localisation: methods and per-channel table

This supplementary block backs Section 4.3.2 (main text). It documents how the per-segment probability → biology correlations are computed and reports the full per-channel Spearman *ρ* table.

#### Per-segment probabilities

Tesseract-VAE (`trans_mean_abmil_v20j_4L`) is evaluated on independent test (*n*_variants_ = 2,950 after the cohort-cleaning filter of Section 4.1) under segmented inference at *S* = 10 (each segment spans 10 windows × 500 bp = 5 kb). The reported score for a segment *s* of variant *v* is the sigmoid output *p*_*v,s*_ ∈ [0, 1] from the classification head on the segment’s *S*-window slice of latents. Segments tile the variant without overlap; a variant of length *L* contributes ⌈*L/*(10 · 500 bp)⌉ segments to the pool.

#### Per-segment biology

For each of the 74 reportable input channels (two channels, `variant_mask` and `length_channel`, are meta annotations excluded from this analysis) we bin the raw bp-resolution track into the same segment grid and apply a kind-appropriate aggregator:

- **continuous peak** (*ρ*, pathogenicity scores): per-segment `max`.
- **continuous density** (histone ChIP, ATAC/DNase): per-segment `mean`.
- **gene-span score** (pLI, LOEUF, HI_index, pHaplo, pTriplo): per-segment `max` over the scored gene spans.
- **ordinal step** (ConsHMM, ChromHMM): per-segment `max` state value.
- **binary span** (ClinGen evidence levels, OMIM, CDS/UTR, ENCODE cCRE categories): per-segment fraction of base pairs *>* 0 (`overlap`).
- **binary point** (ClinVar pathogenic SNV / indel positions): per-segment `count`.
- **sparse continuous peak** (`gnomad_af`): per-segment `max` over non-zero positions.

This produces a single scalar signal *b*_*v,s,c*_ per (variant, segment, channel), pairable 1-to-1 with the probability *p*_*v,s*_. The preferred aggregator per kind is fixed by the schema to avoid cherry-picking; the choice is documented alongside the channel.

#### Cohort-level Spearman *ρ*

For each channel *c* we pool (*p*_*v,s*_, *b*_*v,s,c*_) across every segment of every variant in the independent test (*n*_segments_ = 183,938) and compute the Spearman rank correlation. Because *p*_*v,s*_ is a per-segment sigmoid (no per-variant sum constraint), raw-value pooling is legitimate — there is no within-variant rescaling step of the kind required for attention weights.

#### Cluster-bootstrap 95% CI

Segments from the same variant are dependent — same biological context, adjacent genomic positions, single softmax sampling at training — so a segment-level bootstrap would underestimate variance. We cluster at the variant level: resample *n*_variants_ variants with replacement, each variant carrying all its segments, reconstruct the cohort pool, and recompute *ρ*. Five hundred replicates; the 95% CI is the 2.5th–97.5th percentile of the replicate distribution. Significance at the 5% level is read directly from whether the CI excludes zero.

**Table 14:**
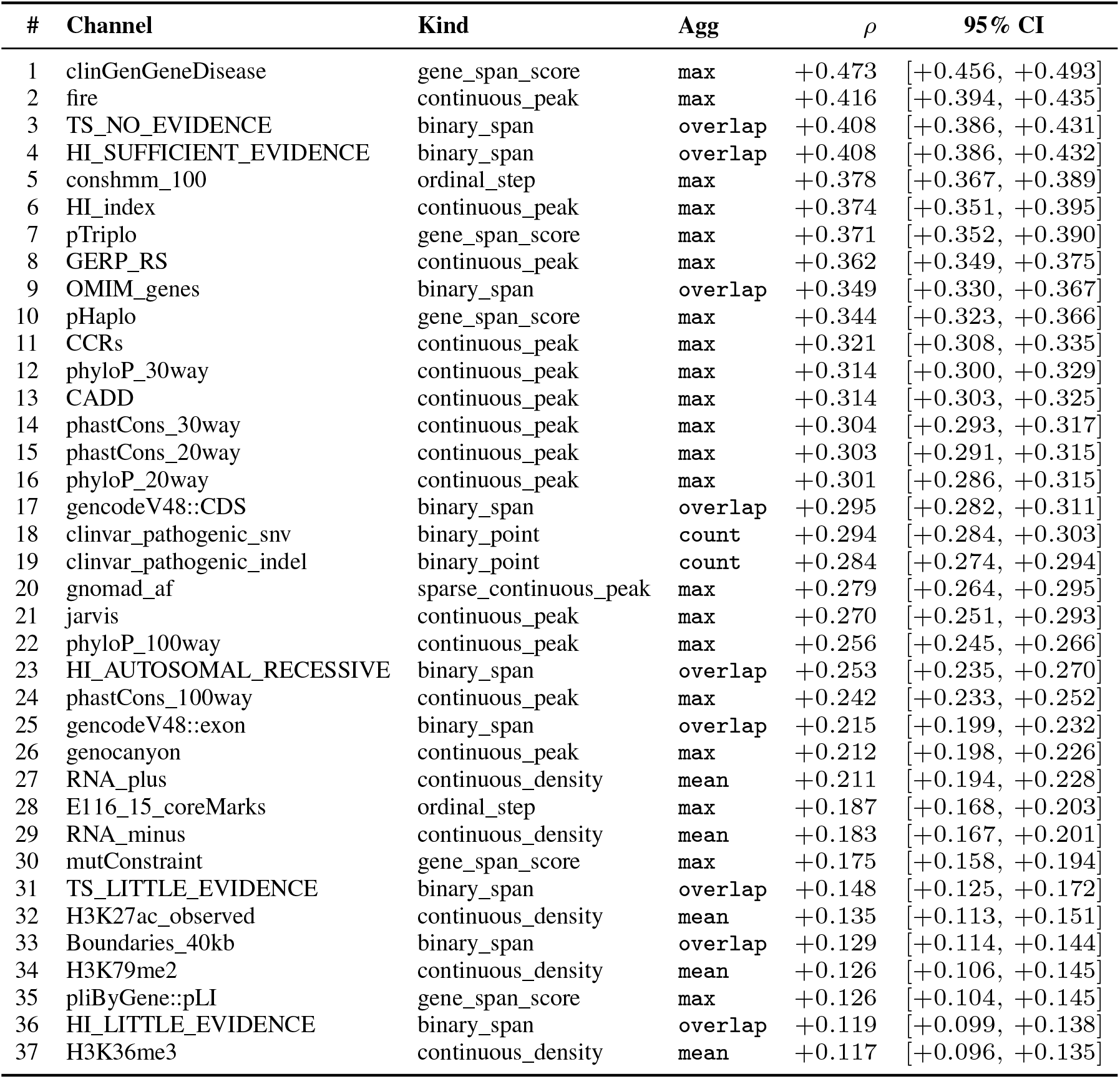
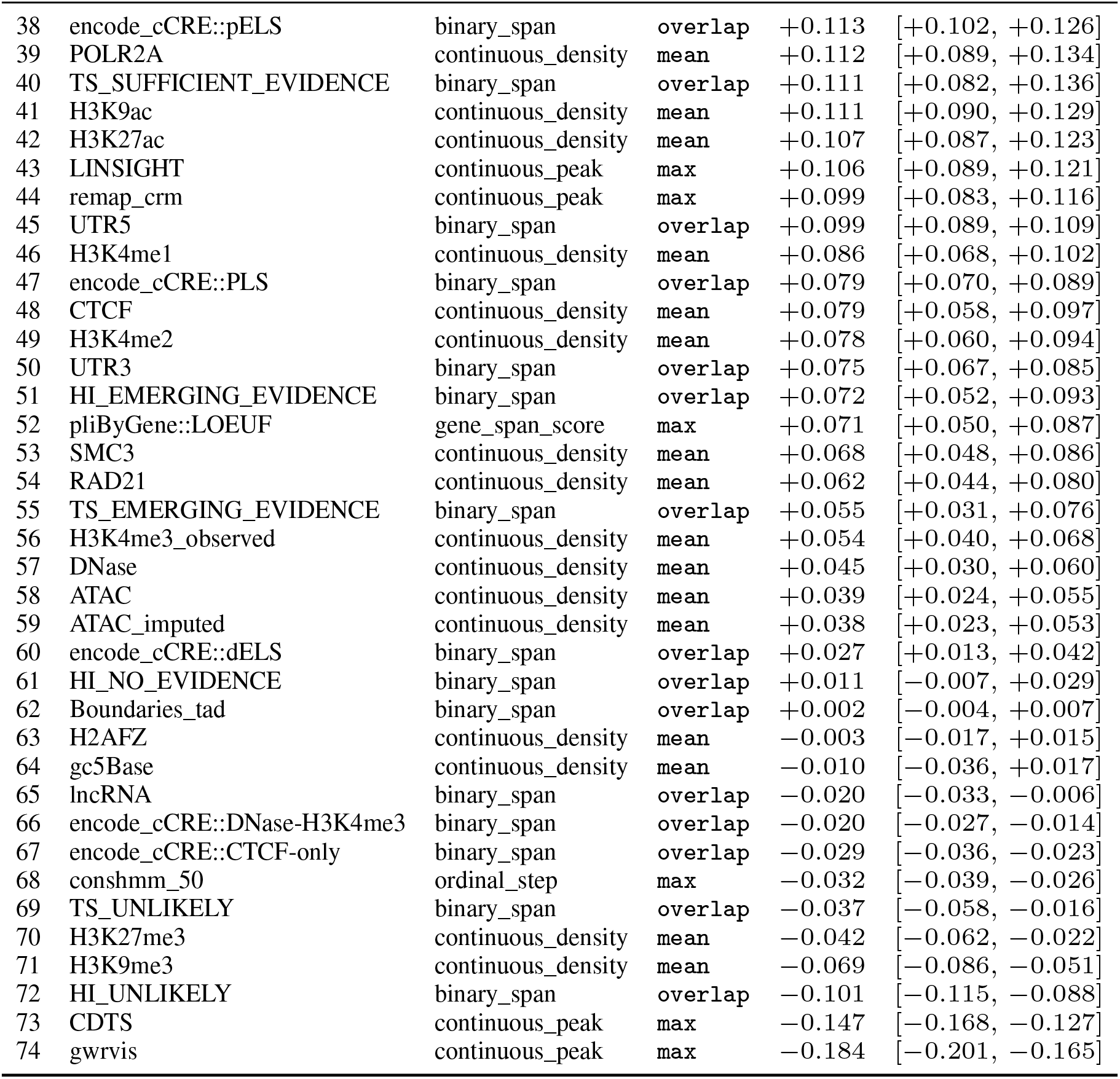
Table S7 — Per-channel Spearman *ρ* (Independent test, segmented inference *S* = 10, Tesseract-VAE). Per-segment probability *p*_*v,s*_ vs per-segment biological-track value under the preferred aggregator per kind (methods above). *n*_variants_ = 2,950; *n*_segments_ = 183,938. CI = 95% cluster-bootstrap (500 replicates, variants as cluster unit). A row is significant at the 5% level iff its CI excludes zero; 70 / 74 rows are significant (60 positive, 10 negative), 4 null. Rows sorted by *ρ*.

#### A.10 Channel perturbation study

We perturb each input-track group by zeroing its tracks at evaluation time and measuring the drop in predictive performance. Both Tesseract-E2E and Tesseract-VAE reuse the pretrained checkpoints from the main benchmark; no retraining is performed. For both models, the selected tracks of the raw (78, *W*) input tensor are set to zero prior to encoding, after which the perturbed input is passed through the otherwise-unchanged model pipeline.

##### Evaluation

Each perturbed configuration is evaluated in full-length inference and compared to the un-perturbed baseline by AUC. Table 15 reports the three cohorts emphasised here—Held-out test coding, Held-out test noncoding, and independent test—showing absolute AUC for the baseline and ΔAUC in percentage points for every perturbation, bolded where |Δ| ≥ 2 pp.

**Table 15:**
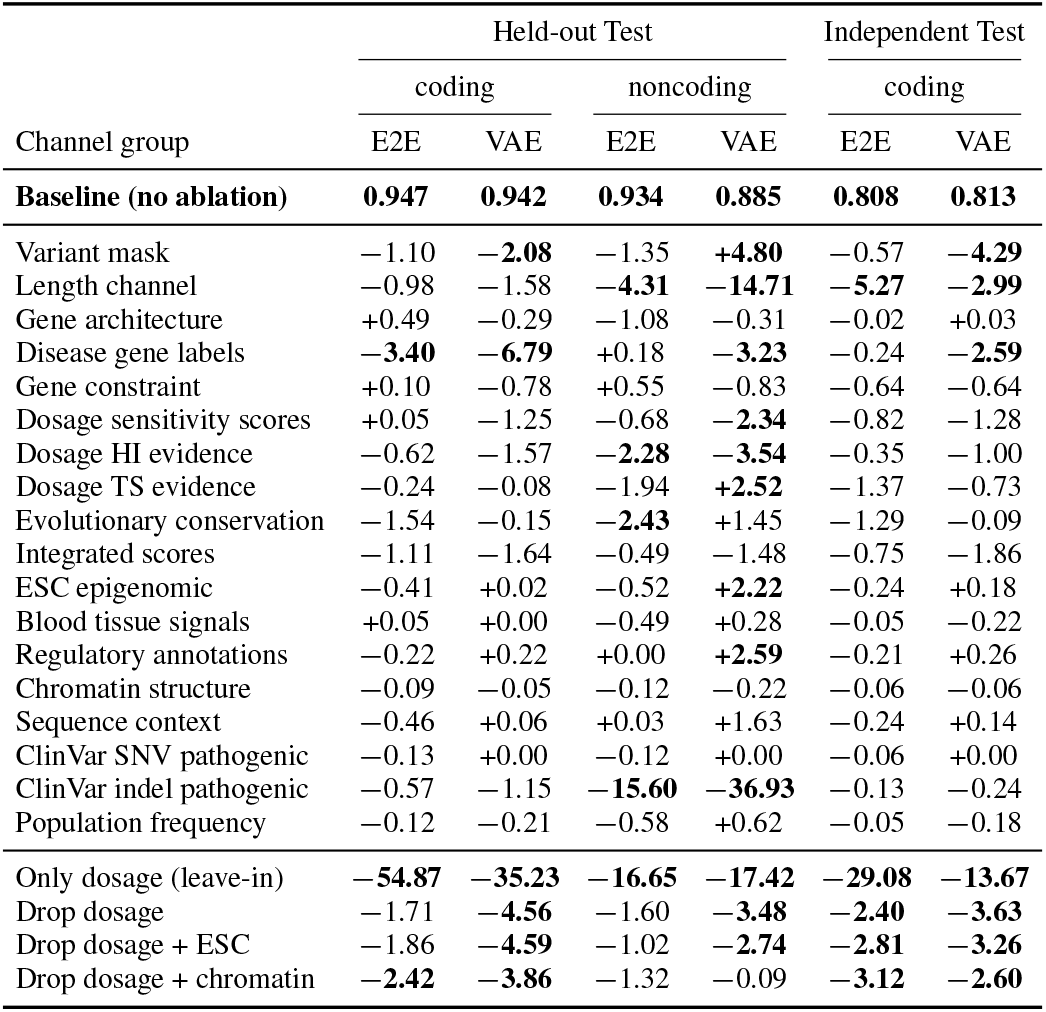
Channel-ablation AUC for TESSERACT-E2E (Raw-CNN) and TESSERACT-VAE (VAE). Baseline row shows absolute AUC. Other rows show ΔAUC in percentage points relative to each model’s own baseline. Entries with |Δ| ≥ 2 pp are bolded. Full-length eval (no segmentation).

| Channel group | Held-out Test |  |  |  | Independent Test |  |
| --- | --- | --- | --- | --- | --- | --- |
|  | coding |  | noncoding |  | coding |  |
|  | E2E | VAE | E2E | VAE | E2E | VAE |
| **Baseline (no ablation)** | **0.947** | **0.942** | **0.934** | **0.885** | **0.808** | **0.813** |
| Variant mask | -1.10 | **-2.08** | -1.35 | **+4.80** | -0.57 | **-4.29** |
| Length channel | -0.98 | -1.58 | **-4.31** | **-14.71** | **-5.27** | **-2.99** |
| Gene architecture | +0.49 | -0.29 | -1.08 | -0.31 | -0.02 | +0.03 |
| Disease gene labels | **-3.40** | **-6.79** | +0.18 | **-3.23** | -0.24 | **-2.59** |
| Gene constraint | +0.10 | -0.78 | +0.55 | -0.83 | -0.64 | -0.64 |
| Dosage sensitivity scores | +0.05 | -1.25 | -0.68 | **-2.34** | -0.82 | -1.28 |
| Dosage HI evidence | -0.62 | -1.57 | **-2.28** | **-3.54** | -0.35 | -1.00 |
| Dosage TS evidence | -0.24 | -0.08 | -1.94 | **+2.52** | -1.37 | -0.73 |
| Evolutionary conservation | -1.54 | -0.15 | **-2.43** | +1.45 | -1.29 | -0.09 |
| Integrated scores | -1.11 | -1.64 | -0.49 | -1.48 | -0.75 | -1.86 |
| ESC epigenomic | -0.41 | +0.02 | -0.52 | **+2.22** | -0.24 | +0.18 |
| Blood tissue signals | +0.05 | +0.00 | -0.49 | +0.28 | -0.05 | -0.22 |
| Regulatory annotations | -0.22 | +0.22 | +0.00 | **+2.59** | -0.21 | +0.26 |
| Chromatin structure | -0.09 | -0.05 | -0.12 | -0.22 | -0.06 | -0.06 |
| Sequence context | -0.46 | +0.06 | +0.03 | +1.63 | -0.24 | +0.14 |
| ClinVar SNV pathogenic | -0.13 | +0.00 | -0.12 | +0.00 | -0.06 | +0.00 |
| ClinVar indel pathogenic | -0.57 | -1.15 | **-15.60** | **-36.93** | -0.13 | -0.24 |
| Population frequency | -0.12 | -0.21 | -0.58 | +0.62 | -0.05 | -0.18 |
| Only dosage (leave-in) | **-54.87** | **-35.23** | **-16.65** | **-17.42** | **-29.08** | **-13.67** |
| Drop dosage | -1.71 | **-4.56** | -1.60 | **-3.48** | **-2.40** | **-3.63** |
| Drop dosage + ESC | -1.86 | **-4.59** | -1.02 | **-2.74** | **-2.81** | **-3.26** |
| Drop dosage + chromatin | **-2.42** | **-3.86** | -1.32 | -0.09 | **-3.12** | **-2.60** |

##### Perturbation groups

The 78 channels are partitioned into 18 biologically coherent *singles*, each isolating one semantic theme:

- **Variant identity**. `variant_mask` (ch 0), `length_channel` (ch 1).
- **Gene architecture**. GENCODE v48 exon / CDS, lncRNA, UTR5, UTR3 (ch 2–6).
- **Disease gene labels**. OMIM, ClinGen gene–disease validity (ch 7, 15).
- **Gene constraint**. pLI, LOEUF, mutConstraint, CCRs (ch 10–13).
- **Dosage sensitivity scores**. pHaplo, pTriplo, HI_index (ch 8, 9, 14).
- **Dosage HI evidence**. ClinGen HI tier tracks (ch 16–21).
- **Dosage TS evidence**. ClinGen TS tier tracks (ch 22–27).
- **Evolutionary conservation**. phyloP, phastCons, GERP_RS, ConsHMM (ch 28–36).
- **Integrated scores**. LINSIGHT, CADD, CDTS, JARVIS, gwRVIS, GenoCanyon, FIRE (ch 37–42, 70).
- **ESC epigenomic**. ESC histone marks and chromatin-accessibility tracks (ch 43–57).
- **Blood tissue tracks**. Blood ATAC, H3K27ac, H3K4me3, RNA ± strand (ch 58–62).
- **Regulatory annotations**. ChromHMM, super-enhancers, ENCODE cCREs, ReMap CRM (ch 63–69, 71).
- **Chromatin structure**. TAD boundaries, 40 kb boundaries (ch 72–73).
- **Sequence context**. GC content (ch 74).
- **ClinVar SNV pathogenic**. (ch 75).
- **ClinVar indel pathogenic**. (ch 76).
- **Population frequency**. gnomAD SNV allele frequency (ch 77).

Singles cover all 78 channels exactly once. We additionally report 5 *compound* perturbations to test cross-group compensation:

- **Only dosage (leave-in)**. Zero everything except the three dosage singles (*dosage prior* + HI + TS evidence, 15 channels kept).
- **Drop dosage**. Zero the three dosage singles (15 channels).
- **Drop dosage + ESC**. Drop dosage plus ESC epigenomic (30 channels).
- **Drop dosage + chromatin**. Drop dosage plus the full chromatin stack: ESC epigenomic, blood tissue signals, regulatory annotations, chromatin architecture (45 channels).

##### Broader impact

Tesseract could improve rare-disease CNV interpretation by prioritizing pathogenicity-relevant genomic subregions and making model predictions more spatially inspectable by clinical genetics researchers. Potential positive impacts include improved triage of candidate CNVs, more efficient review of large structural variants, and better prioritization of genomic evidence for downstream expert interpretation.

The main risks arise from over-reliance on automated pathogenicity scores in clinical settings. Incorrect predictions could contribute to misclassification of variants, inappropriate prioritization of evidence, or unequal performance across genomic regions, variant classes, or patient populations that are underrepresented in curated CNV databases. Because the model is trained on existing clinical labels and annotation tracks, it may also inherit biases from ascertainment, reporting practices, and uneven annotation quality. Tesseract is therefore intended as a research and evidence-prioritization tool, not as a standalone diagnostic system. Clinical use would require external validation, calibration, prospective evaluation, and interpretation by qualified clinical genetics professionals.

